# Quantifying the effect of experimental perturbations at single-cell resolution

**DOI:** 10.1101/532846

**Authors:** Daniel B. Burkhardt, Jay S. Stanley, Alexander Tong, Ana Luisa Perdigoto, Scott A. Gigante, Kevan C. Herold, Guy Wolf, Antonio J. Giraldez, David van Dijk, Smita Krishnaswamy

**Author notes:** These authors contributed equally (Daniel B. Burkhardt, Jay S. Stanley). These authors contributed equally (Guy Wolf, Antonio J. Giraldez, David van Dijk, Smita Krishnaswamy).

## Abstract

Current methods for comparing scRNA-seq datasets collected in multiple conditions focus on discrete regions of the transcriptional state space, such as clusters of cells. Here, we quantify the effects of perturbations at the single-cell level using a continuous measure of the effect of a perturbation across the transcriptomic space. We describe this space as a manifold and develop a relative likelihood estimate of observing each cell in each of the experimental conditions using graph signal processing. This likelihood estimate can be used to identify cell populations specifically affected by a perturbation. We also develop vertex frequency clustering to extract populations of affected cells at the level of granularity that matches the perturbation response. The accuracy of our algorithm to identify clusters of cells that are enriched or depleted in each condition is on average 57% higher than the next best-performing algorithm tested. Gene signatures derived from these clusters are more accurate compared to six alternative algorithms in ground-truth comparisons.

## 1 Introduction

As single-cell RNA-sequencing (scRNA-seq) has become more accessible, the design of single-cell experiments has become increasingly complex. Researchers regularly use scRNA-seq to quantify the effect of a drug, gene knockout, or other experimental perturbation on a biological system. However, quantifying the differences between single-cell datasets collected from multiple experimental conditions remains an analytical challenge [1]. This task is hindered by biological heterogeneity, technical noise, and uneven exposure to a perturbation. Furthermore, each single-cell dataset comprises several intrinsic structures of heterogeneous cells, and the effect of the treatment condition could be diffuse across all cells or isolated to particular populations. To address this, we develop a method that quantifies the probability that each cell state would be observed in a given sample condition.

Our goal is to quantify the effect of an experimental perturbation on every cell observed in matched treatment and control scRNA-seq samples of the same biological system. We begin by modelling the cellular transcriptomic state space as a smooth low-dimensional manifold or set of manifolds. This approach has been previously applied to characterize cellular heterogeneity and dynamic biological processes in single cell data [2–8]. We then define and calculate a *sample-associated density estimate,* which quantifies the density of each sample over the manifold of cell states. We then consider differences in the sample-associated density estimates for each cell to calculate a *sample-associated relative likelihood,* which quantifies the effect of an experimental perturbation as the likelihood of observing each cell in each experimental condition (**Figure 1**).

**Figure 1:**
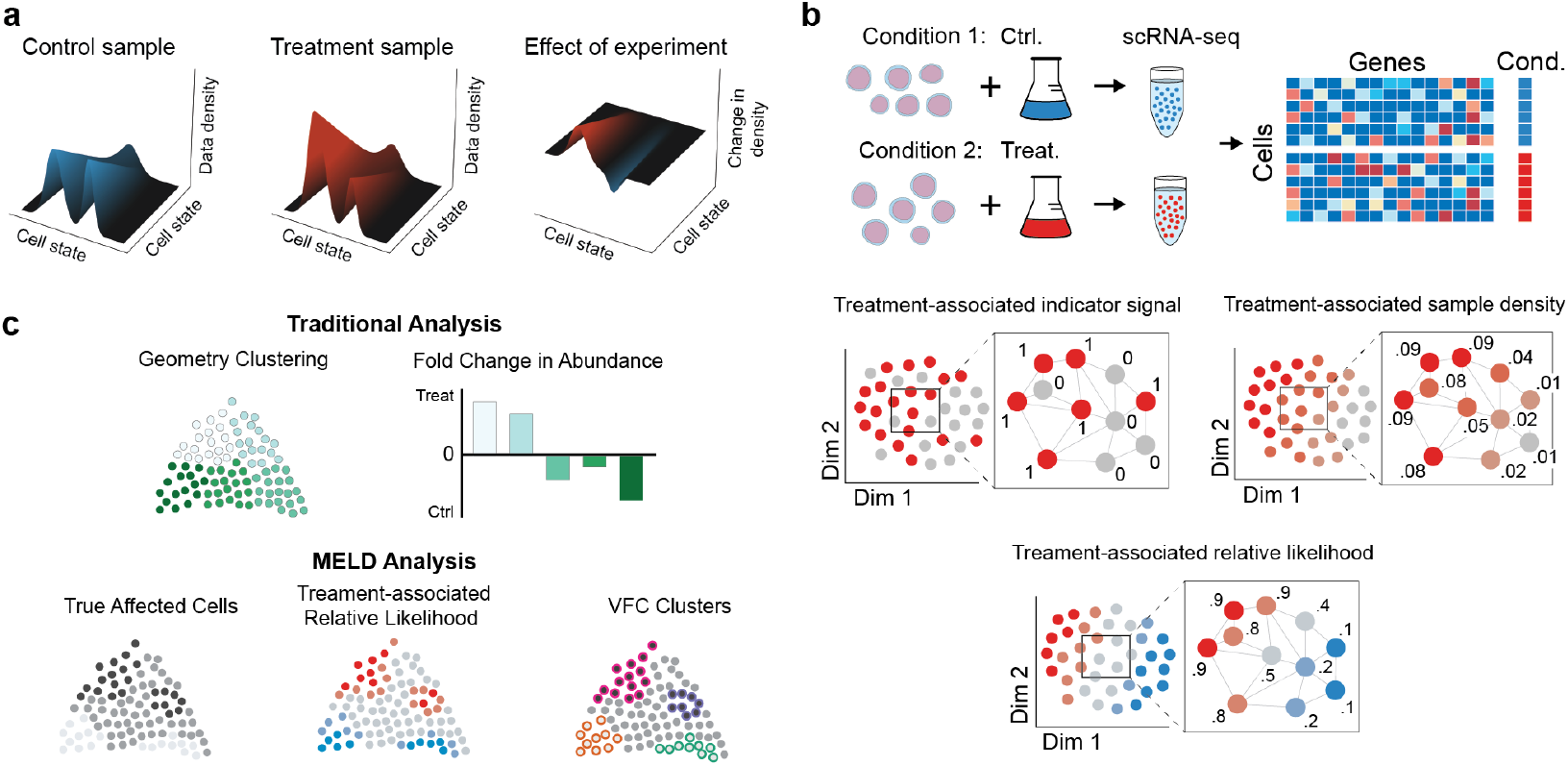
(**a**) To quantify the effect of an experiment, we model single cell experiments as samples from a probability density function (pdf) over the underlying transcriptomic cell state space manifold. The pdf for the control sample is the frequency with which cell states are observed in the control sample compared to the overall frequency of the cell state in both samples combined. In this context, the effect of an experimental perturbation is to alter this probability density and thus the data density in the treatment sample relative to the control. Therefore, the effect of an experimental perturbation can be quantified as the change in the probability density in the experiment condition relative to the control. (**b**) The sample-associated relative likelihood quantifies this effect by computing a kernel density estimate over the cell similarity graph using graph signals representing indicator vectors for each sample. The sample-associated relative likelihood indicates the likelihood that a particular cell is from the treatment or control conditions. (**c**) In traditional analysis of scRNA-seq datasets, the clusters are based solely on the data geometry and changes in abundance between conditions may not align with the true affected populations. Using the sample-associated relative likelihood and VFC, we can identify the correct cluster resolution for downstream analysis.

Almost all previous work quantifying differences between single cell datasets relies on discrete partitioning of the data prior to downstream analysis [9–16]. First, datasets are merged applying either batch normalization [15, 16] or a simple concatenation of data matrices [9–14]. Next, clusters are identified by grouping either sets of cells or modules of genes. Finally, within each cluster, the cells from each condition are used to calculate statistical measures, such as fold-change between samples. However, reducing experimental analysis to the level of clusters sacrifices the power of single-cell data. We demonstrate cases where subsets of a cluster exhibit divergent responses to a perturbation that were missed in published analysis that was limited to clusters derived using data geometry alone. Instead of quantifying the effect of a perturbation within clusters, we focus on the level of single cells.

In the sections that follow, we show that the sample-associated relative likelihood has useful information for the analysis of experimental conditions in scRNA-seq. First, the relative likelihoods of each condition can be used to identify the cell states most and least affected by an experimental treatment. Second, we show that the frequency composition of the sample label and the relative likelihood scores can be used as the basis for a clustering algorithm we call *vertex frequency clustering* (VFC). VFC identifies populations of cells that are similarly affected (either enriched, depleted, or unchanged) between conditions at the level of granularity of the perturbation response. Third, we obtain gene signatures of a perturbation by performing differential expression between vertex frequency clusters.

We call the algorithm to calculate the sample-associated density estimate and relative likelihood the MELD algorithm, so named for its utility in joint analysis of single-cell datasets. The MELD and VFC algorithms are provided in an open-source Python package available on GitHub at https://github.com/KrishnaswamyLab/MELD.

## 2 Results

### 2.1 Overview of the MELD algorithm

We propose a framework for quantifying differences in cell states observed across single-cell samples. The power of scRNA-seq as a measure of an experimental treatment is that it provides samples of cell state at thousands to millions of points across the transcriptomic space in varying experimental conditions. Our approach is inspired by recent successes in applying manifold learning to scRNA-seq analysis [17]. The manifold model is a useful approximation for the transcriptomic space because biologically valid cellular states are intrinsically low-dimensional with smooth transitions between similar states. In this context, our goal is to quantify the change in enrichment of cell states along the underlying cellular manifold as a result of the experimental treatment (**Figure 1**).

For an intuitive understanding, we first consider a simple experiment with one sample from a treatment condition and one sample from a control condition. Here, sample refers to a library of scRNA-seq profiles, and condition refers to a particular configuration of experimental variables. In this simple experiment, our goal is to calculate the relative likelihood that each cell would be observed in either the treatment or control condition over a manifold approximated from all cells from both conditions. This relative likelihood can be used as a measure of the effect of the experimental perturbation because it indicates for each cell how much more likely we are to observe that cell state in the treatment condition relative to the control condition (**Figure 1**). We refer to this ratio as the *sample-associated relative likelihood.* The steps to calculate the sample-associated relative likelihood are given in **Algorithm 1** and a visual depiction can be found in **Figure S1.**

As has been done previously, we first approximate the cellular manifold by constructing an affinity graph between cells from all samples [2–8]. In this graph, each node corresponds to a cell, and the edges between nodes describe the transcriptional similarity between the cells. We then estimate the density of each sample over the graph using graph signal processing [18]. A graph signal is any function that has a defined value for each node in a graph. Here we use labels indicating the sample origin of each cell to develop a collection of one-hot indicator signals over the graph with one signal per sample. Each indicator signal has value 1 associated with each cell from the corresponding sample and value 0 elsewhere. In a simple two-sample experiment, the sample indicator signals would comprise two one-hot signals, one for the control sample and one for the treatment sample. These one-hot signals are column-wise L1 normalized to account for different numbers of cells sequenced in each sample. After normalization, each indicator signal represents an empirical probability density over the graph for the corresponding sample. We next use these normalized indicator signals to calculate a kernel density estimate of each sample over the graph.

#### Algorithm 1

The MELD algorithm

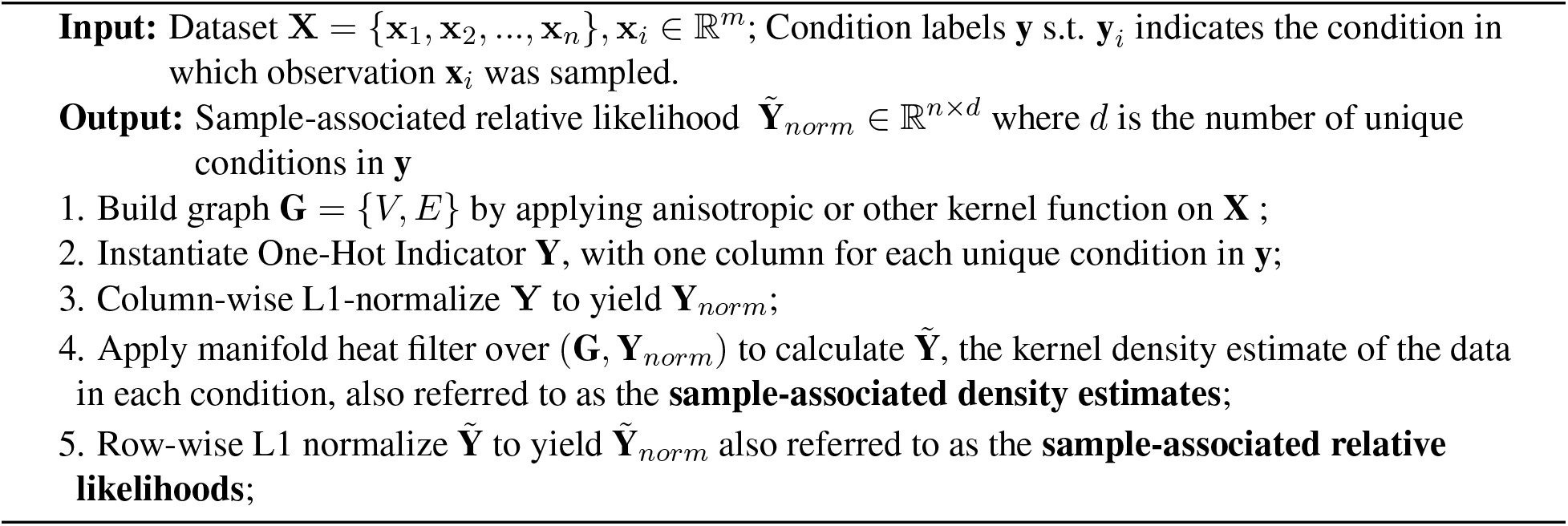

### 2.2 Calculating sample-associated density estimates

A popular non-parametric approach to estimating data density is using a kernel density estimate (KDE), which relies on an affinity kernel function. To estimate the density of single cell samples over a graph, we turn to the heat kernel. This kernel uses diffusion to provide local adaptivity in regions of varying data density [19] such as is observed in single cell data. Here, we extend this kernel as a low pass filter over a graph to estimate the density of a sample represented by the sample indicator signals defined above. To begin, we take the Gaussian KDE, which is a well known tool for density estimation in ℝ^*d*^. We then generalize this form to smooth manifolds. The full construction of this generalization is described in detail in **Section 4.1.10**, and a high level overview is provided here.

A kernel density estimator 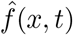 with bandwidth *t* > 0 and kernel function *K*(*x, y, t*) is defined as

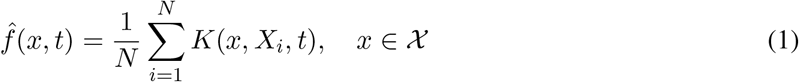

where *X* is the observed data, *x* is some point in 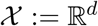 (i.e., 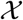 is defined as ℝ^*d*^), and 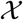 is endowed with the Gaussian kernel defined as

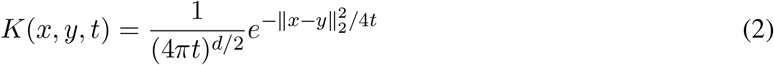

Thus, **Equation 2** defines the Gaussian KDE in ℝ^*d*^. However, this function relies on the Euclidean distance 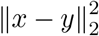, which is derived from the kernel space in ℝ^*d*^. Since manifolds are only locally Euclidean, we cannot apply this KDE directly to a general manifold.

To generalize the Gaussian KDE to a manifold we need to define a kernel space (i.e., the range of a kernel operator) over a manifold. In ℝ^*d*^ the kernel space is often defined via infinite weighted sums of sines and cosines, also known as the Fourier series. However, this basis is not well defined for a Riemannian manifold, so we instead use the eigenbasis of the Laplace operator as our kernel basis. The derivation and implication of this extension is formally explored in **Section 4.1.10**. The key insight is that using this kernel space, the Gaussian KDE can be defined as a filter constructed from the eigenvectors and eigenvalues of the Laplace operator on a manifold. When this manifold is approximated using a graph, we define this KDE as a graph filter over the graph Laplacian given by the following equation:

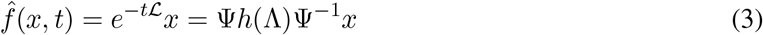

where *t* is the kernel bandwidth, 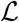 is the graph Laplacian, *x* is the empirical density, Ψ and Λ are the eigenvectors and corresponding eigenvalues of 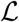, and 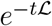 is the matrix exponential. This signal processing formulation can alternatively be formulated as an optimization with Tikhonov Regularization, which seeks to reconstruct the original signal while penalizing differences along edges of the graph. This connection is further explored in **Section 4.1.7**.

To achieve an efficient implementation of the filter in **Equation 3**, the MELD algorithm considers the spectral representation of the sample indicator signals and uses a Chebyshev polynomial approximation [20] to efficiently compute the sample-associated density estimate (see **Section 4.1.4**). The result is a highly scalable implementation. The sample-associated density estimate for two conditions can be calculated on a dataset of 50,000 cells in less than 8 minutes in a free Google Colaboratory notebook^1^, with more than 7 minutes of that time spent constructing a graph that can be reused for visualization [3] or imputation [4]. With the sample-associated density estimates, it is now possible to identify the cells that are most and least affected by an experimental perturbation.

### 2.3 Using sample-associated relative likelihood to quantify differences between experimental conditions

Each sample-associated density estimate over the graph indicates the probability of observing each cell within a given experimental sample. For example, in a healthy peripheral blood sample, we would expect high density estimates associated with abundant blood cells such as neutrophils and T cells and low density estimates associated with less abundant cells types, such as basophils and eosinophils. When considering the effect of an experimental perturbation, we are not only interested in these density estimates directly, but we want also to quantify the change in density associated with a change in an experimental variable. For example, one might want to know if a drug treatment causes a change in probability of observing some kinds of blood cells in peripheral blood.

When examining the rows of the sample-associated density estimates for a single cell, the values represent the likelihood of observing that cell in each experimental condition. To quantify the change in likelihood across conditions, we apply a normalization across the likelihoods for each cell to calculate sample-associated relative likelihoods. These relative likelihoods sum to 1 for each cell and provide a basis for quantifying the change in likelihood of observing a cell in each condition. We then use these relative likelihoods as a basis for identifying cell states that are enriched, depleted, or unaffected by the perturbation.

The sample-associated relative likelihoods can be used to analyze scRNA-seq perturbation studies of varying experimental designs. For cases with only one experimental and one control condition, we typically only refer to the sample-associated relative likelihood of the treatment condition for downstream analysis. For more complicated experiments comprising replicates, we normalize matched experimental and control conditions individually, then average the relative likelihood of the each condition across replicates, as in **Section 2.7** and **Section 2.8**. With datasets comprising three or more experimental conditions, each sample-associated relative likelihood may be used individually to analyze cells that are enriched, depleted, or unaffected in the corresponding condition, as in **Section 2.9**. We expect this flexibility will enable the use of sample-associated density estimates and relative likelihoods across a wide range of single cell studies.

### 2.4 Vertex-frequency clustering identifies cell populations affected by a perturbation

A common goal for analysis of experimental scRNA-seq data is to identify subpopulations of cells that are responsive to the experimental treatment. Existing methods cluster cells by transcriptome alone and then attempt to quantify the degree to which these clusters are differentially represented in the two conditions. However, this is problematic because the granularity, or sizes, of these clusters may not correspond to the sizes of the cell populations that respond similarly to experimental treatment. Additionally, when partitioning data along a continuum, cluster boundaries are somewhat arbitrary and may not correspond to populations with distinct differences between conditions. Our goal is to identify clusters that are not only transcriptionally similar but also respond similarly to an experimental perturbation (**Figure 2**).

**Figure 2:**
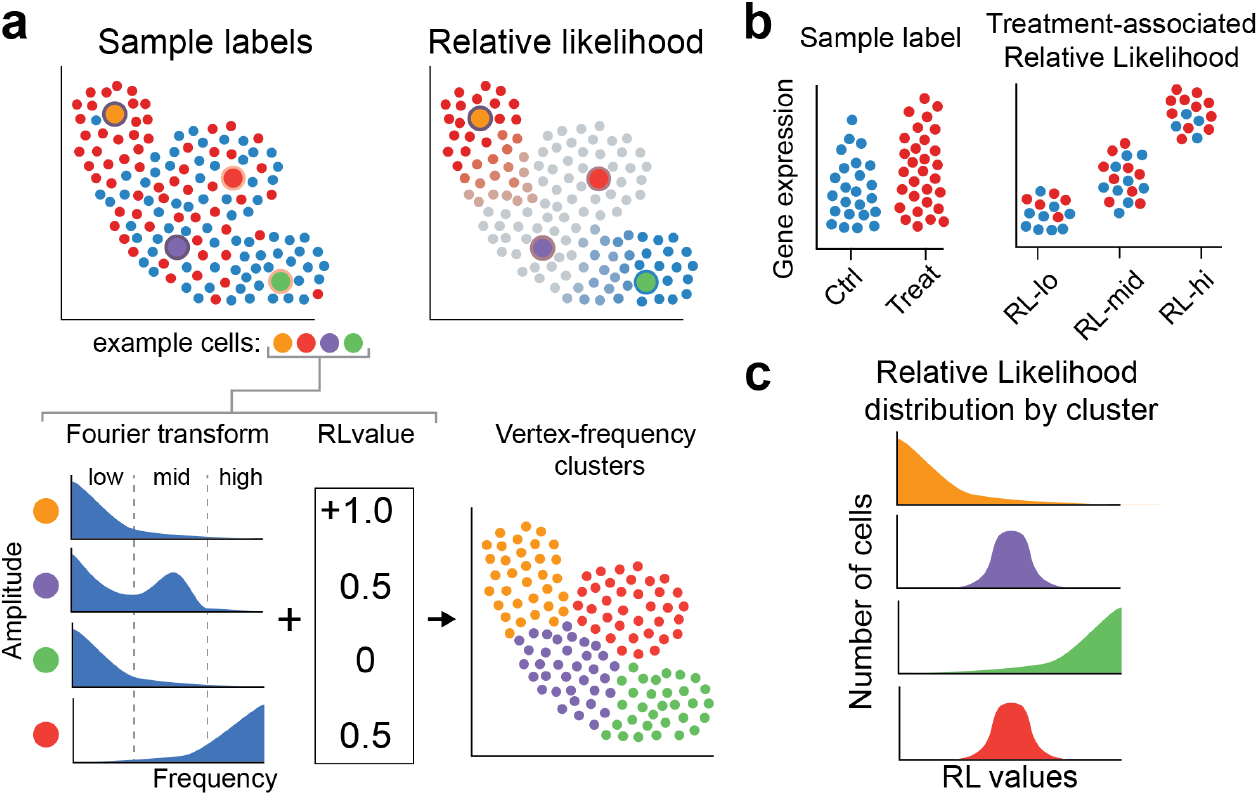
Vertex Frequency Analysis using the sample-associated indicator signals and relative likelihood (**a**) The Windowed Graph Fourier Transform of the sample-associated indicator signals and values of sample-associated relative likelihood values at four example points shows distinct patterns between a transitional (blue) and unaffected (red) cell. This information is used in spectral clustering, resulting in Vertex Frequency Clustering. (**b**) Characterizing Vertex Frequency Clusters with the highest and lowest sample-associated relative likelihood values elucidates gene expression changes associated with experimental perturbations. (**c**) Examining the distribution of sample-associated relative likelihood scores in vertex-frequency clusters identifies cell populations most affected by a perturbation.

A naïve approach to identify such clusters would be to simply concatenate the sample-associated relative likelihood to the gene expression data as an additional feature and cluster on these combined features. However, the magnitude of the relative likelihood does not give a complete picture of differences in response to a perturbation. For example, even in a two-sample experiment, there are multiple ways for a cell to have a sample-associated relative likelihood of 0.5. In one case, it might be that there is a continuum of cells one end of which is enriched in the treatment condition and the other end is enriched in the control condition. In this case transitional cells halfway through this continuum will have a sample-associated relative likelihood of 0.5 (we show an example of this in **Section 2.6**). Another scenario that would result in a relative likelihood of 0.5 is even mixing of a population of cells between control and treatment conditions with no transition, i.e., cells that are part of a non-responsive cell subtype that is unchanged between conditions (we show an example of this in **Section 2.8** and **Figure S2**). To differentiate between such scenarios we must consider not only the magnitude of the sample-associated relative likelihood but also the frequency of the input sample indicator signals over the manifold. Indeed in the transitional case the input sample labels change gradually or has *low frequency* over the manifold, and in the even-mixture case it changes frequently between closely connected cells or has *high frequency* over the manifold.

As no contemporary method is suitable for resolving these cases, we developed an algorithm that integrates gene expression, the magnitude of sample-associated relative likelihoods, and the frequency response of the input sample labels over the cellular manifold (**Figure S2**). In particular, we cluster using local frequency profiles of the sample indicator signal around each cell. This method, which we call *vertex-frequency clustering* (VFC), is an adaptation of the signal-biased spectral clustering proposed by Shuman et al. [21]. The VFC algorithm provides a feature basis for clustering based on the spectrogram [21] of the sample indicator signals, which can be thought of as a histogram of frequency components of graph signals. We observe that we can distinguish between non-responsive populations of cells with high frequency sample indicator signal components and transitional populations with lower frequency indicator signal components. The VFC feature basis combines this frequency information with the magnitude of the sample-associated relative likelihood and the cell similarity graph to identify phenotypically similar populations of cells with uniform response to a perturbation. The algorithm is discussed in further detail in **Section 4.2**.

With VFC, it is possible to define a new paradigm for recovering the gene signature of a perturbation. In traditional analysis, where clusters are calculating data geometry alone, gene signatures are often calculated using differential expression analysis between experimental conditions within each cluster (**Figure S3a**). The theory of the traditional framework is that these expression differences reflect the change in cell states observed as a result of the perturbation. However, if the cluster contains multiple subpopulations that each contain different responses to the perturbation, we can first separate these populations using VFC and then compare each subpopulation individually (**Figure S3b**). Not only does this allow for more finely resolved comparisons, we show in the following section that this approach is capable of recovering gene signatures more accurately than directly comparing two samples.

We describe a full pipeline for analysis of scRNA-seq datasets with MELD and VFC in **Supplementary Note 7.1** and **Figure S4**.

### 2.5 Quantitative validation of the MELD and VFC algorithms

No previous benchmarks exist to quantify the ability of an algorithm to capture changes in density between scRNA-seq samples. To validate the sample-associated relative likelihood and VFC algorithms, we used a combination of simulated scRNA-seq data and synthetic experiments using previously published datasets. To create simulated scRNA-seq data, we used Splatter [22]. To ensure the algorithms worked on real scRNA-seq datasets, we also used two previously published datasets comprising Jurkat T cells [13] and cells from whole zebrafish embryos [15]. In each dataset, we created a ground truth relative likelihood distribution over all cells that determines the relative likelihood each cell would be observed in one of two simulated conditions. In each simulation, different populations of cells of varying sizes were depleted or enriched. Cells were then randomly split into two samples according to this ground truth relative likelihood and used as input to each algorithm. More detail on the comparison experiments is provided in **Section 4.7**.

We performed three sets of quantitative comparisons. First, we calculated the degree to which the MELD algorithm captured the ground truth relative likelihood distribution in each simulation. We found that MELD outperformed other graph smoothing algorithms by 10-52% on simulated data and 36-51% on real datasets (**Figure 3 Table S1**). We also determined that the MELD algorithm is robust to the number of cells captured in the experiment with only a 10% decrease in performance when 65% of the cells in the T cell dataset were removed (**Figure S5**). We used results from these simulations to determine the optimal parameters for the MELD algorithm (**Section 4.3**). Next, we quantified the accuracy of the VFC algorithm to identify clusters of cells that were enriched or depleted in each condition. When compared to six common clustering algorithms including Leiden [23] and CellHarmony [24], VFC was the top performing algorithm on every simulation on the T cell data and best performing on average on the zebrafish dataset with a 57% increase in average performance over Louvain, the next best algorithm (**Figures S6a-c & S7**, **Table S2**). Finally, we calculated how well VFC clusters could be used to calculate the gene signature of a perturbation. Gene signatures obtained using VFC were compared to signatures obtained using direct comparison of two conditions–the current standard–and those obtained using other clustering algorithms (**Figure S6d**). These results confirm that MELD and VFC outperform existing methods for analyzing multiple scRNA-seq datasets from different experimental conditions.

**Figure 3:**
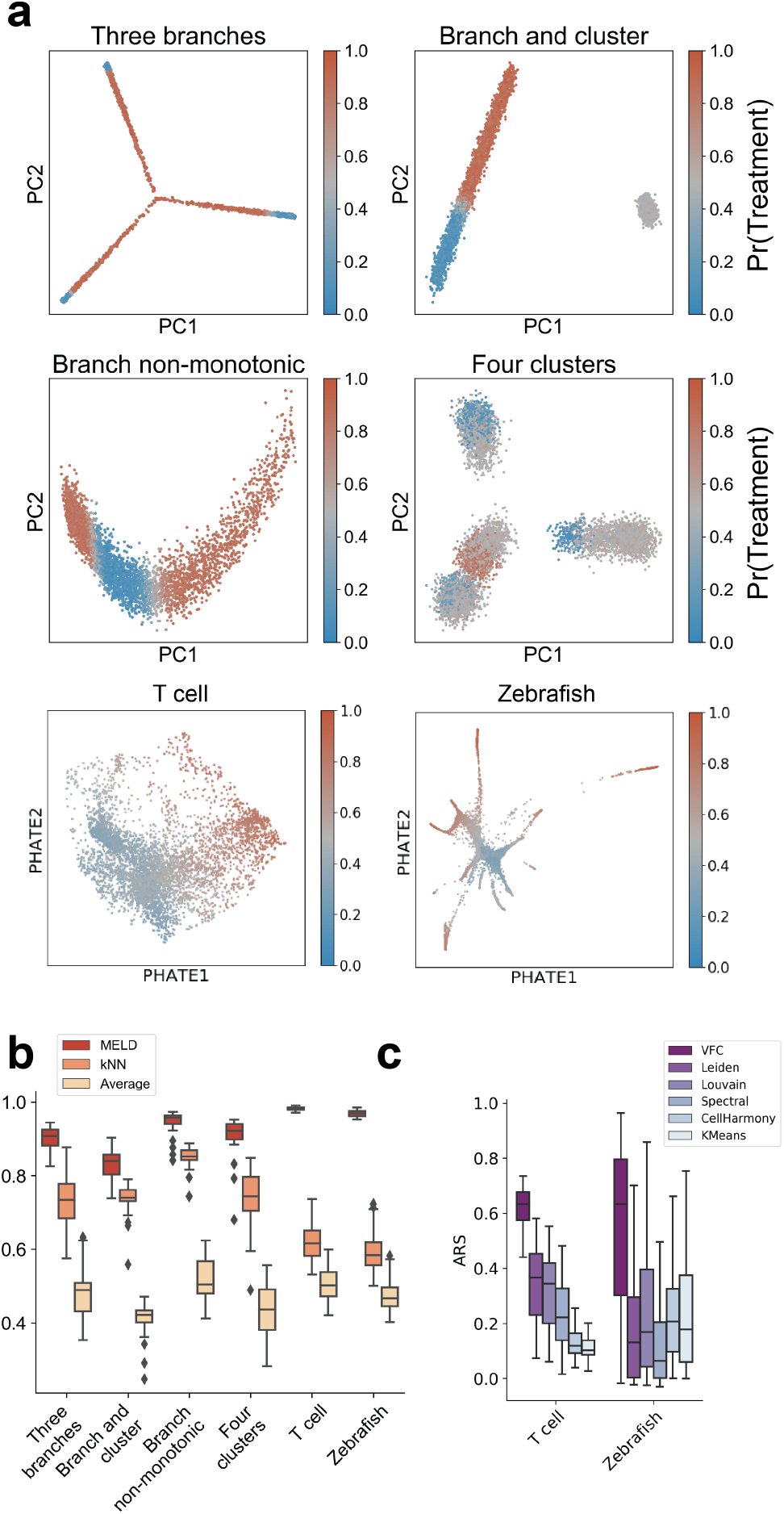
Quantitative comparison of the sample-associated Relative Likelihood and VFC. (**a**) Single cell datasets were generated using Splatter [22] or taken from previously published experiments [13, 15]. Ground truth sample assignment probabilities with each of two conditions were randomly generated 20 times with varying noise and regions of enrichment for the simulated data and 100 random sample assignments were generated for the real-world datasets. Each cell is colored by the probability of being assigned to the treatment sample. (**b**) Pearson correlation comparison of the sample-associated relative likelihood algorithm to kNN averaging of the sample labels and graph averaging of the sample labels. Higher values are better. (**c**) Comparison of VFC to popular clustering algorithms. Adjusted Rand Score (ARS) quantifies how accurately each method detects regions that were enriched, depleted, or unchanged in the experimental condition relative to the control. Higher values are better.

### 2.6 The sample-associated relative likelihood identifies a biologically relevant signature of T cell activation

To demonstrate the biological relevance of the MELD algorithm, we analyze Jurkat T cells cultured for 10 days with and without anti-CD3/anti-CD28 antibodies as part of a Cas9 knock-out screen published by Datlinger et al. [13] (**Figure 4a**). The goal of this experiment was to characterize the transcriptional signature of T cell Receptor (TCR) activation and determine the impact of gene knockouts in the TCR pathway. First, we visualized cells using PHATE, a visualization and dimensionality reduction tool for single-cell RNA-seq data (**Figure 4b**) [3]. We observed a large degree of overlap in cell states between the stimulated and control conditions, as noted in the original study [13].

**Figure 4:**
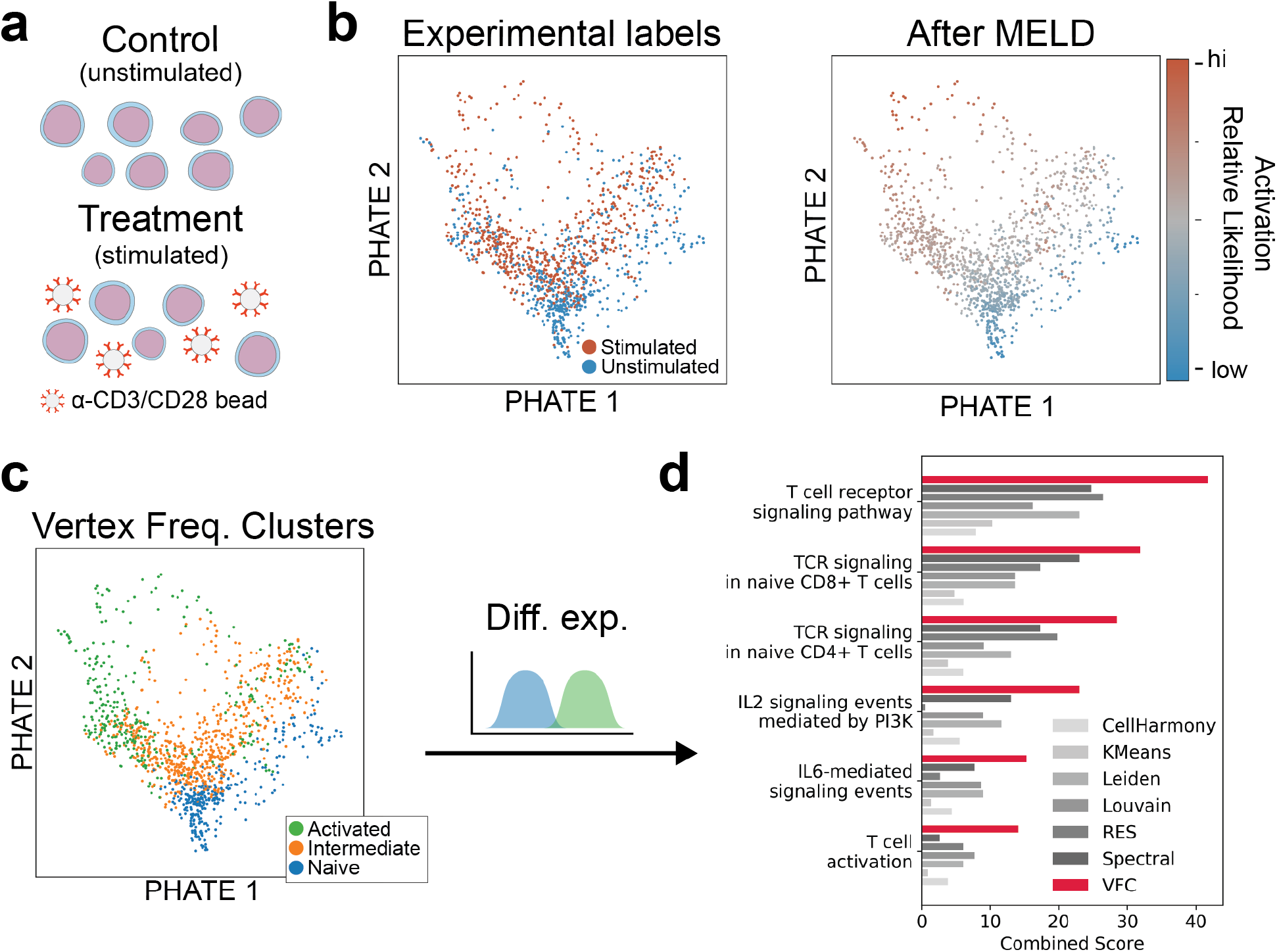
MELD recovers signature of TCR activation. (**a**) Jurkat T-cells were stimulated with α-CD3/CD28 coated beads for 10 days before collection for scRNA-seq. (**b**) Examining a PHATE plot, there is a large degree of overlap in cell state between experimental conditions. However, after MELD it is clear which cells states are prototypical of each experimental condition. (**c**) Vertex Frequency Clustering identifies an activated, a naive, and an intermediate population of cells. (**d**) Signature genes identified by comparing the activated to naive cells are enriched for annotations related to TCR activation using EnrichR analysis. Combined scores for the MELD gene signature are shown in red and scores for a gene signature obtained using the sample labels only are shown in grey.

To determine a gene signature of the TCR activation, we considered cells with no CRISPR perturbation. First, we computed sample-associated relative likelihood and VFC clusters on these samples. Then we derived a gene signature by performing differential expression analysis between VFC clusters with the highest and lowest relative likelihood values. We identified 2335 genes with a q-value < 0.05 as measured by a rank sum test with a Benjamini & Hochberg False Discovery Rate correction [25]. We then compared this signature to those obtained using the same methods from our simulation experiments. To determine the biological relevance of these signature genes, we performed gene set enrichment analysis on both gene sets using EnrichR [26]. Considering the GO terms highlighted by Datlinger et al. [13], we found that the MELD gene list has the highest combined score in all of the gene terms we examined (**Figure 4d**). These results show that the sample-associated relative likelihood and VFC are capable of identifying a biologically relevant dimension of T cell activation at the resolution of single cells. Furthermore, the gene signature identified using the MELD and VFC outperformed standard differential expression analyses to identify the signature of a real-world experimental perturbation.

Finally, to quantitatively rank the impact of each Cas9 gene knockout on TCR activation we examined the distribution of sample-associated relative likelihood values for all stimulated cells transfected with gRNAs targeting a given gene (**Figure S8**). We observed a large variation in the impact of each gene knockout consistent with the published results from Datlinger et al. [13]. Encouragingly, our results agree with the bulk RNA-seq validation experiment of Datlinger et al. [13] showing strongest depletion of TCR response with knockout of kinases LCK and ZAP70 and adaptor protein LAT. We also find a slight increase in relative likelihood of the stimulation condition in cells in which negative regulators of TCR activation are knocked out, including PTPN6, PTPN11, and EGR3. Together, these results show that the MELD and VFC algorithms are suitable for characterizing a biological process such as TCR activation in the context of a complex Cas9 knockout screen.

### 2.7 VFC improves characterization of subpopulation response to *chd* loss-of-function

To demonstrate the utility of sample-associated relative likelihood analysis applied to datasets composed of multiple cell types, we analyzed a chordin loss-of-function experiment in zebrafish using CRISPR/Cas9 (**Figure 5**) [15]. In the experiment published by Wagner et al. [15], zebrafish embryos were injected at the 1-cell stage with Cas9 and gRNAs targeting either chordin (*chd*), a BMP-antagonist required for developmental patterning, or tyrosinase (*tyr*), a control gene. Embryos were collected for scRNA-seq at 14-16 hours post-fertilization (hpf). We expect incomplete penetrance of the perturbation in this dataset because of the mosaic nature of Cas9 mutagenesis [27].

**Figure 5:**
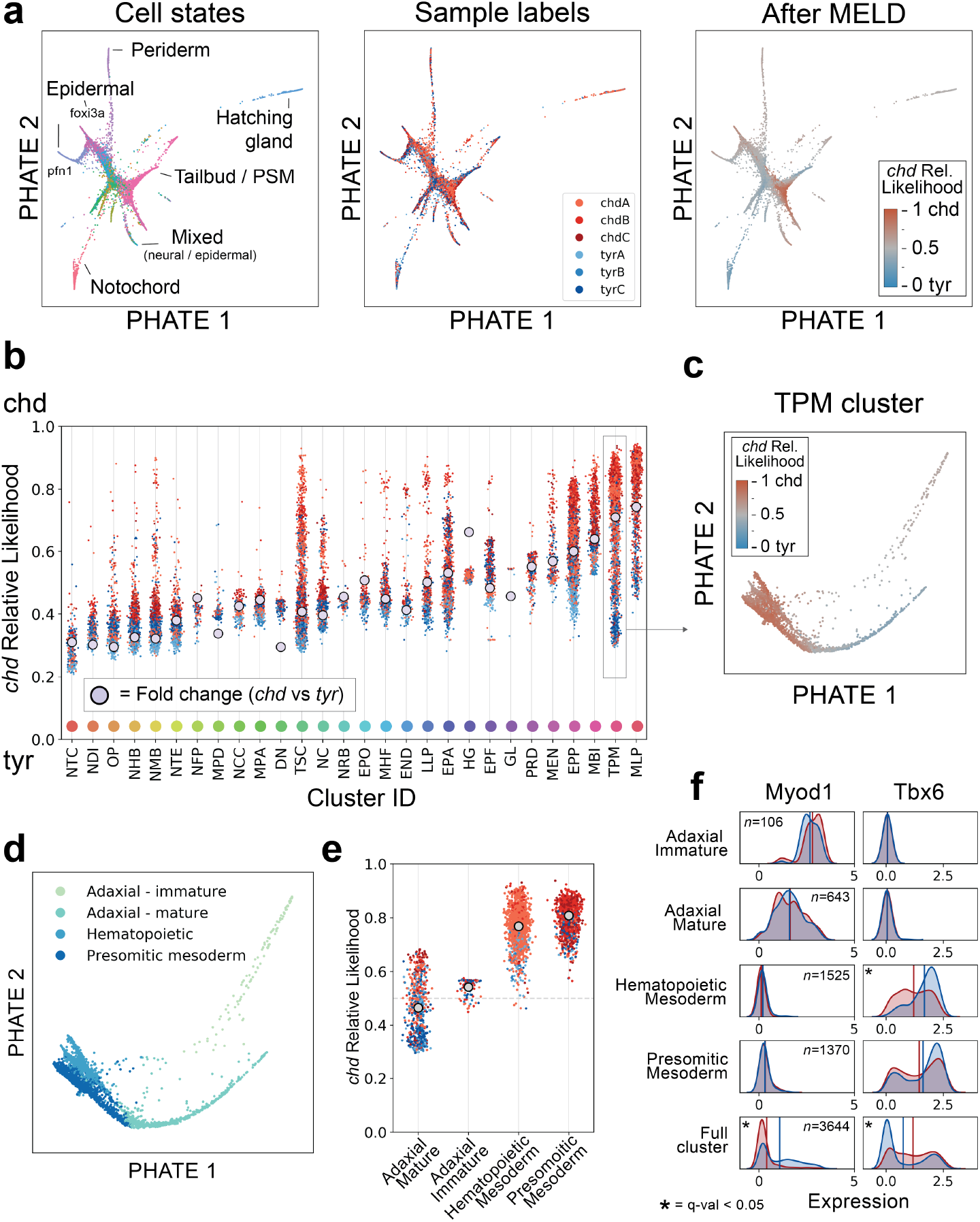
Characterizing chordin Cas9 mutagenesis with MELD. (**a**) PHATE shows a high degree of overlap of sample labels across cell types. Applying MELD to the mutagenesis vector reveals regions of cell states enriched in the *chd* or *tyr* conditions. (**b**) Using published cluster assignments^2^, we show that the *chd*-associated relative likelihood quantifies the effect of the experimental perturbation on each cell, providing more information than calculating fold-change in the number of cells between conditions in each cluster (grey dot), as was done in the published analysis. Color of each point corresponds to the sample labels in panel (a). Generally, average relative likelihood within each cluster aligns with the fold-change metric. However, we can identify clusters, such as the TPM or TSC, with large ranges of relative likelihoods indicating non-uniform response to the perturbation. (**c**) Visualizing the TPM cluster using PHATE, we observe several cell states with mostly non-overlapping relative likelihood values. (**d**) Vertex Frequency Clustering identifies four cell types in the TPM. (**e**) We see the range of relative likelihood values in the TPM cluster is due to subpopulations with divergent responses to the *chd* perturbation. (**f**) We observe that changes in gene expression between the *tyr* (blue) and *chd* (red) conditions is driven mostly by changes in abundance of subpopulations with the TPM cluster.

First, we calculate the sample-associated relative likelihood between the chordin and tyrosinase conditions. Because the experiment was performed in triplicate with three paired *chd* and *tyr* samples, we first calculated the sample-associated density estimates for each of the six samples. We then normalized the density estimated across the paired *chd* and *tyr* conditions. Finally, we averaged the replicate-specific relative likelihoods of the *chd* condition for downstream analysis. We refer to this averaged likelihood simply as the chordin relative likelihood (**Figure S9**).

To characterize the effect of mutagenesis on various cell populations, we first examined the distribution of chordin relative likelihood values across the 28 cell state clusters generated by Wagner et al. [15] (**Figure 5b**). We find that overall the most enriched clusters contain mesodermal cells and the most depleted clusters contain dorsally-derived neural cells matching the ventralization phenotype previously reported with *chd* loss-of-function [28–30]. However, we observe that several clusters have a wide range of chordin relative likelihood values suggesting that there are cells in these clusters with different perturbation responses. Using VFC analysis we find that several of these clusters contain biologically distinct subpopulations of cells with divergent responses to *chd* knock out.

An advantage of using MELD and VFC is the ability to characterize the response to the perturbation at the resolution corresponding to the perturbation response (**Figure 2c**). We infer that the resolution of the published clusters is too coarse because the distribution of chordin relative likelihood values is very large for several of the clusters. For example the chordin relative likelihoods within the Tailbud – Presomitic Mesoderm (TPM) range from 0.29-0.94 indicating some cells are strongly enriched while others are depleted. To disentangle these effects, we performed VFC subclustering for all clusters using the strategy proposed in **Section 7.1**. We found 12 of the 28 published clusters warranted further subclustering with VFC resulting in a total of 50 final cluster labels (**Figure S10j**). To determine the biological relevance of the VFC clusters, we manually annotated each of the three largest clusters subdivided by VFC revealing previously unreported effects of *chd* loss-of-function within this dataset. A full exploration can be found in **Supplementary Note 7.2** with the results of TPM cluster shown in **Figure 5c-f**.

### 2.8 Identifying the effect of IFN*γ* stimulation on pancreatic islet cells

To determine the ability of the MELD and VFC to uncover biological insights, we generated and characterized a dataset of human pancreatic islet cells cultured for 24 hours with and without interferon-gamma (IFN*γ*), a system with significant clinical relevance to auto-immune diseases of the pancreas such as Type I Diabetes mellitus (T1D) and islet allograft rejection [31]. Previous studies have characterized the effect of these cytokines on pancreatic beta cells using bulk RNA-sequencing[32], but no studies have addressed this system at single-cell resolution.

To better understand the effect of immune cytokines on islet cells, we cultured islet cells from three donors for 24 hours with and without IFN*γ* and collected cells for scRNA-seq. After filtering, we obtained 5,708 cells for further analysis. Examining the expression of marker genes for major cell types of the pancreas, we observed a noticeable batch effect associated with the donor ID, driven by the maximum expression of glucagon, insulin, and somatostatin in alpha, beta, and delta cells respectively (**Figure S11a**). To correct for this difference while preserving the relevant differences between donors, we applied the MNN kernel correction described in Section 4.1.1. Note, here we are applying the MNN correction is only applied across donors, not across the IFNg treatment. We developed guidelines for applying batch correction prior to running MELD in **Supplementary Note 7.3**.

To quantify the effect of IFN*γ* treatment across these cell types, we calculated the sample-associated relative likelihood of IFN*γ* stimulation using the same strategy to handle matched replicates as was done for the zebrafish data (**Figure 6a**). We then used established marker genes of islet cells [33] to identify three major populations of cells corresponding to alpha, beta, and delta cells (**Figures 6a-b & S11b**). We next applied VFC to each of the three endocrine cell types and identified a total of nine clusters. Notably, we found two clusters of beta cells with intermediate IFNg relative likelihood values. These clusters are cleanly separated on the PHATE plot of all islet cells (**Figure 6a**) and together the beta cells represent the largest range of IFNg relative likelihood scores in the dataset.

**Figure 6:**
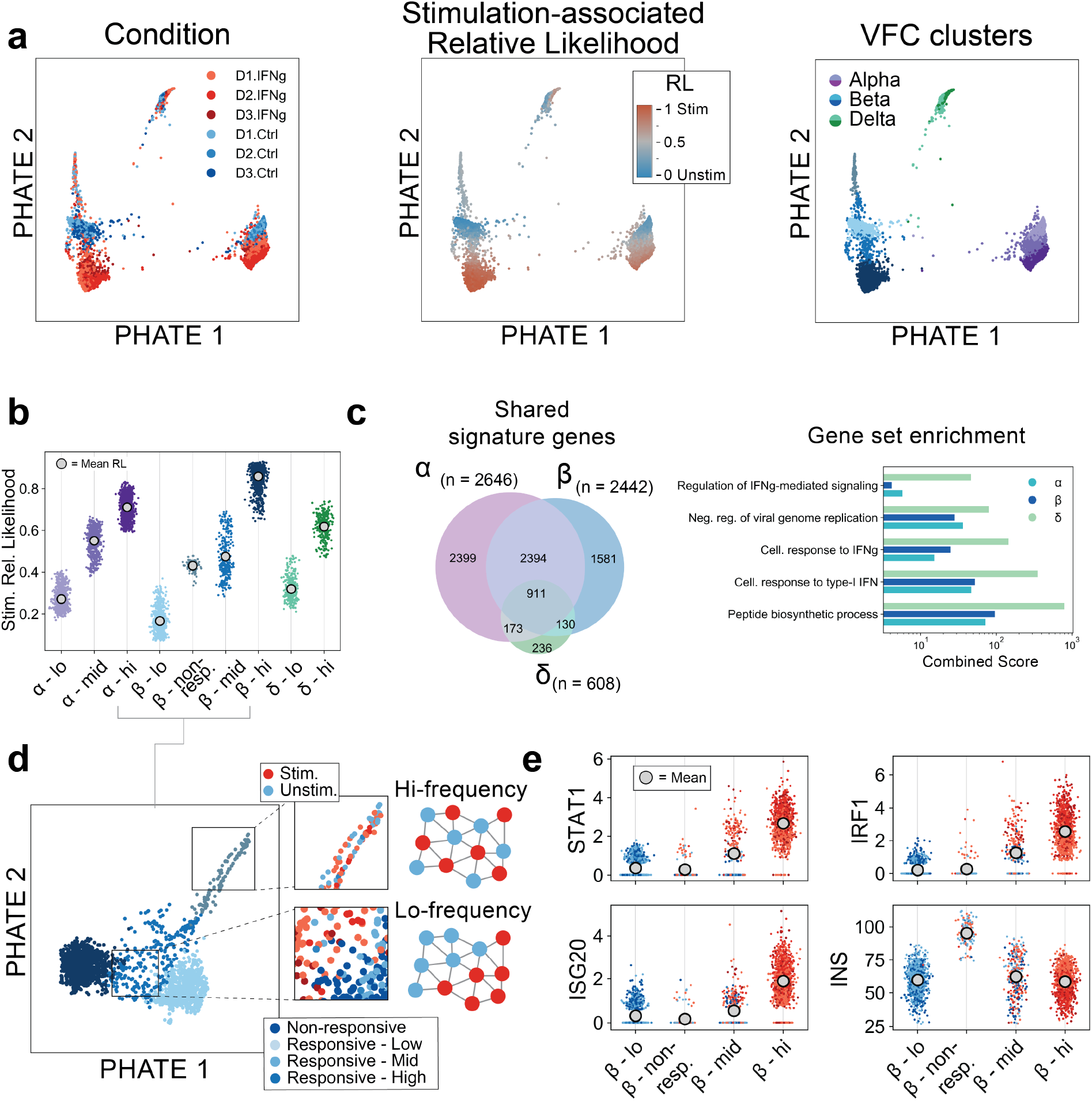
MELD characterizes the response to IFN*γ* in pancreatic islet cells. (**a**) PHATE visualization of pancreatic islet cells cultured for 24 hours with or without IFN*γ*. Vertex-frequency clustering identifies nine clusters corresponding to alpha, beta, and delta cells. (**b**) Examining the stimulation-associated relative likelihood (RL) in each cluster, we observe that beta cells have a wider range of responses than alpha or delta cells. (**c**) We identify the signature of IFN*γ* stimulation by calculating differential expression between the VFC clusters with the highest and lowest stimulation likelihood values for each cell type. We find a high degree of overlap of the significantly differentially expressed genes between alpha and beta cells. (**d**) Results of gene set enrichment analysis for signature genes in each cell type. Beta cells have the strongest enrichment for IFN response pathway genes. (**e**) Examining the four beta cell clusters more closely, we observe two populations with intermediate relative likelihood values. These populations are differentiated by the structure of the sample label in each cluster (outset). In the non-responsive cluster, the sample label has very high frequency unlike the low frequency pattern in the transitional Responsive – mid cluster. (**f**) We find that the non-responsive cluster has low expression of IFN*γ*-regulated genes such as STAT1 despite containing roughly equal numbers of unstimulated and stimulated cells. This cluster is marked by approximately 40% higher expression of insulin.

To further inspect these beta cell clusters, we consider a separate PHATE plot of the cells in the four beta cell clusters (**Figure 6e**). Examining the distribution of input sample signals values in these intermediate cell types, we find that one cluster, which we label as *Non-responsive*, exhibits high frequency input sample signals indicative of a population of cells that does not respond to an experimental treatment. The *Responsive – Mid* cluster matches our characterization of a transitional population with a structured distribution of input sample signals. Supporting this characterization, we find a lack of upregulation in IFN*γ*-regulated genes such as STAT1 in the non-responsive cluster, similar to the cluster of beta cells with the lowest IFNg relative likelihood values (**Figure 6f**).

In order to understand the difference between the non-responsive beta cells and the responsive populations, we calculated differential expression of genes in the non-responsive clusters and all others as previously described [4]. The gene with the greatest difference in expression was insulin, the major hormone produced by beta cells, which is approximately 2.5-fold increased in the non-responsive cells (**Figure 6f**). This cluster of cells bears resemblance to a recently described “extreme” population of beta cells that exhibit elevated insulin mRNA levels and are found to be more abundant in diabetic mice[34, 35]. That these cells appear non-responsive to IFN*γ* stimulation and exhibit extreme expression of insulin suggests that the presence of extreme high insulin in a beta cell prior to IFN*γ* exposure may inhibit the IFN*γ* response pathway through an unknown mechanism.

We next characterized the gene expression signature of IFN*γ* treatment across all three endocrine cell types (**Figure 6c-d**). Using a rank sum test to identify genes that change the most between the clusters with highest and lowest IFNg relative likelihood values within each endocrine population, we identify 911 genes differentially expressed in all three cell types. This consensus signature includes activation of genes in the JAK-STAT pathway including STAT1 and IRF1 [36] and in the IFN-mediated antiviral response including MX1, OAS3, ISG20, and RSAD2 [37–39]. The activation of both of these pathways has been previously reported in beta cells in response to IFN*γ* [40, 41]. To confirm the validity of our gene signatures, we use EnrichR [26] to perform gene set enrichment analysis on the signature genes and find strong enrichment for terms associated with interferon signalling pathways (**Figure S11d**). From these results we conclude that although IFN*γ* leads to upregulation of the canonical signalling pathways in all three cell types, the response to stimulation in delta cells is subtly different to that of alpha or beta cells.

Here, we applied MELD analysis to identify the signature of IFN*γ* stimulation across alpha, beta, and delta cells, and we identified a population of beta cells with high insulin expression that appears unaffected by IFN*γ* stimulation. Together, these results demonstrate the utility of MELD analysis to reveal biological insights in a clinically-relevant biological experiment.

### 2.9 Analysis of donor-specific composition

Although most of the analysis here focuses on two condition experiments, we show that it is possible to use the sample-associated relative likelihood to quantify the differences between more than two conditions. In the islet dataset, we have samples of treatment and control scRNA-seq data from three different donors. To quantify the differences in cell profiles between donors, we first create a one-hot vector for each donor label and normalize across all three smoothed vectors. This produces a measure of how likely each transcriptional profile is to be observed in donor 1, 2, or 3. We then analyze each of these signals for each cluster examined in **Section 2.8** (**Figure S12**). We find that all of the alpha cell and delta cell clusters are depleted in donor 3 and the non-responsive beta cell cluster is enriched primarily in donor 1. Furthermore, the most highly activated alpha cell cluster is enriched in donor 2. As with the sample-associated relative likelihood derived for the IFN*γ* response, it is also possible to identify donor-specific changes in gene expression, or clusters of cells differentially abundant between each donor. We propose that this strategy could be used to extend MELD analysis to experiments with multiple categorical experimental conditions, such as data collected from different tissues or stimulus conditions.

## 3 Discussion

When performing multiple scRNA-seq experiments in various experimental and control conditions, researchers often seek to characterize the cell types or sets of genes that change from one condition to another. However, quantifying these differences is challenging due to the subtlety of most biological effects relative to the biological and technical noise inherent to single-cell data. To overcome this hurdle, we designed the MELD and VFC algorithms to quantify compositional differences between samples. The key innovation in the sample-associated relative likelihood algorithm is quantifying the effect of a perturbation at the resolution of single cells using theory from manifold learning.

We have shown that our analysis framework improves over the current best-practice of clustering cells based on gene expression and calculating differential abundance and differential expression within clusters. Clustering prior to quantifying compositional differences can fail to identify the divergent responses of subpopulations of cells within a cluster. Using the sample labels and sample-associated relative likelihood, we apply VFC to derive clusters of cells to identify cells that are most enriched in either condition and cells that are unaffected by an experimental perturbation. We show that gene signatures extracted using these clusters outperform those derived from direct comparison of two samples.

We demonstrated the application of MELD analysis on single-cell datasets from three different biological systems and experimental designs. We provided a framework for handling paired experimental and control replicates and guidance on analysis of complex experimental designs with more than two conditions and in the context of a single-cell Cas9 knockout screen. In our analysis of the zebrafish dataset, we showed the published clusters contained biologically relevant subpopulations of cells with divergent responses to the experimental perturbation. We also described a previously unpublished dataset of pancreatic islet cells stimulated with IFN-γ and characterize a previously unreported subpopulation of β cells that appeared unresponsive to stimulation. We related this to emerging research describing a β cells subtype marked by high insulin mRNA expression and unique biological responses.

We anticipate MELD to have widespread use in many contexts since experimental labels can arise in many contexts. As we showed, if we have sets of single cell data from healthy individuals vs sick individuals, the sample-associated relative likelihood could indicate cell types specific to disease. This framework could potentially be extended to patient level measurements where patients’ phenotypes as measured with clinical variables and laboratory values can be associated with enriched states in disease or treatment conditions. Indeed MELD has already seen use in several contexts [42–46].

## 4 Methods

In this section, we will provide details about our computational methods for computing the sample-associated density estimate and relative likelihood, as well as extracting information from the sample label and sample-associated relative likelihood by way of a method we call *vertex frequency clustering*. We will outline the mathematical foundations for each algorithm, explain how they relate to previous works in manifold learning and graph signal processing, and provide details of the implementations of each algorithm.

### 4.1 Computation of the sample-associated density estimate

Computing the sample-associated density estimate and relative likelihood involves the following steps each of which we will describe in detail.

1. A cell similarity graph is built over the combined data from all samples where each node or vertex in the graph is a cell and edges in the graph connect cells with similar gene expression values.
2. The sample label for each cell is used to create the sample-associated indicator signal.
3. Each indicator signal is then smoothed over the graph to estimate the density of each sample using the manifold heat filter.
4. Sample-associated density estimates for paired treatment and control samples are normalized to calculate the sample-associated relative likelihood.

#### 4.1.1 Graph construction

The first step in the MELD algorithm is to create a cell similarity graph. In single-cell RNA sequencing, each cell is measured as a vector of gene expression counts measured as unique molecules of mRNA. Following best practices for scRNA-seq analysis [1], we normalize these counts by the total number of Unique Molecular Indicators (UMIs) per cell to give relative abundance of each gene and apply a squareroot transform. Next we compute the similarity all pairs of cells, by using their Euclidean distances as an input to a kernel function. More formally, we compute a similarity matrix *W* such that each entry *W_ij_* encodes the similarity between cell gene expression vectors x_i_ and x_j_ from the dataset *X*.

In our implementation we use *α*-decaying kernel proposed by Moon et al. [3] because in practice it provides an effective graph construction for scRNA-seq analysis. However, in cases where batch, density, and technical artifacts confound graph construction, we also use a mutual nearest neighbor kernel as proposed by Haghverdi et al. [47].

The *α*-decaying kernel [3] is defined as

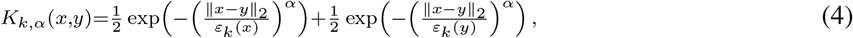

where *x, y* are data points, *ε_k_*(*x*), *ε_k_*(*y*) are the distance from *x, y* to their *k*-th nearest neighbors, respectively, and *α* is a parameter that controls the decay rate (i.e., heaviness of the tails) of the kernel. This construction generalizes the popular Gaussian kernel, which is typically used in manifold learning, but also has some disadvantages alleviated by the *α*-decaying kernel, as explained in Moon et al. [3].

The similarity matrix effectively defines a weighted and fully connected graph between cells such that every two cells are connected and that the connection between cells *x* and *y* is given by *K*(*x,y*). To allow for computational efficiency, we sparsify the graph by setting very small edge weights to 0.

While the kernel in **Equation 4** provides an effective way of capturing neighborhood structure in data, it is susceptible to batch effects. For example, when data is collected from multiple patients, subjects, or environments (generally referred to as “batches”), such batch effects can cause affinities within each batch are often much higher than between batches, thus artificially creating separation between them rather than follow the underlying biological state. To alleviate such effects, we adjust the kernel construction using an approach inspired by recent work from by Haghverdi et al. [47] on the Mutual Nearest Neighbors (MNN) kernel. We extend the standard MNN approach, which has previous been applied to the k-Nearest Neighbors kernel, to the *α*-decay kernel as follows. First, within each batch, the affinities are computed using **Equation 4**. Then, across batches, we compute slightly modified affinities as

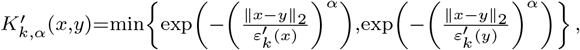

where 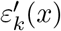 are now computed via the *k*-th nearest neighbor of *x* in the batch containing *y* (and vice versa for 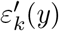). Next, a rescaling factor *γ_xy_* is computed such that

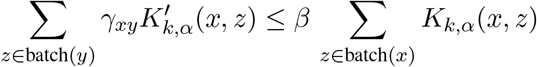

for every *x* and *y*, where *β* > 0 is a user configurable parameter. This factor gives rise to the rescaled kernel

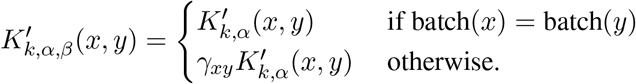

Finally, the full symmetric kernel is then computed as

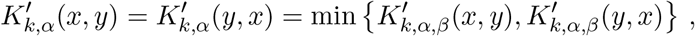

and used to set the weight matrix for the constructed graph over the data. Note that this construction is a well-defined extension of (**Equation 4**), as it reduces back to that kernel when only a single batch exists in the data.

We also perform an anisotropic density normalization transformation so that the kernel reflects the underlying geometry normalized by density as in Coifman and Lafon [48]. The density normalized kernel 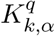 divides out by density, estimated by the sum of outgoing edge weights for each node is as follows,

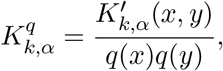

where

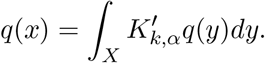

We use this density normalized kernel in all experiments. When the data is uniformly sampled from the manifold then the density around each point is constant then this normalization has no effect. When the density is non-uniformly sampled from the manifold this allows an estimation of the underlying geometry unbiased by density. This is especially important when performing density estimation from empirical distributions with different underlying densities. By normalizing by density, we allow for construction of the manifold geometry from multiple differently distributed samples and individual density estimation for each of these densities on the same support. This normalization is further discussed in **Section 4.1.10**.

#### 4.1.2 Estimating sample-associated density and relative likelihood on a graph

Density estimation is difficult in high dimensions because the number of samples needed to accurately reconstruct density with bounded error is exponential in the number of dimensions. Since general high dimensional density estimation is an intrinsically difficult problem, additional assumptions must be made. A common assumption is that the data exists on a manifold of low intrinsic dimensionality in ambient space. Under this assumption a number of works on graphs have addressed density estimation limited to the support of the graph nodes [49–53]. Instead of estimating kernel density or histograms in *D* dimensions where *D* could be large, these methods rendered the data as a graph, and density is estimated each point on the graph (each data point) as some variant counting the number of points which lie within a radius of each point on the graph.

The MELD algorithm also estimates density of a signal on a graph. We use a generalization of the standard heat kernel on the graph to estimate density (See **Section 4.1.7**). We draw analogs between the resulting sample-associated density estimate and Gaussian kernel density estimation on the manifold showing our density estimate with a specific parameter set is equivalent to the Gaussian density estimate on the graph (See **Section 4.1.10**).

#### 4.1.3 Graph Signal Processing

The MELD algorithm leverages recent advances in graph signal processing (GSP) [18], which aim to extend traditional signal processing tools from the spatiotemporal domain to the graph domain. Such extensions include, for example, wavelet transforms [54], windowed Fourier transforms [21], and uncertainty principles [55]. All of these extensions rely heavily on the fundamental analogy between classical Fourier transform and graph Fourier transform (described in the next section) derived from eigenfunctions of the graph Laplacian, which is defined as

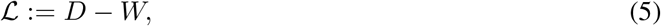

where *D* is the *degree* matrix, which is a diagonal matrix with *D_ii_* = *d*(*i*) = ∑_*j*_ *W_ij_* containing the degrees of the vertices of the graph defined by *W*.

#### 4.1.4 The Graph Fourier Transform

One of the fundamental tools in traditional signal processing is the Fourier transform, which extracts the frequency content of spatiotemporal signals [56]. Frequency information enables various insights into important characteristics of analyzed signals, such as pitch in audio signals or edges and textures in images. Common to all of these is the relation between frequency and notions of *smoothness*. Intuitively, a function is *smooth* if one is unlikely to encounter a dramatic change in value across neighboring points. A simple way to imagine this is to look at the *zero-crossings* of a function. Consider, for example, sine waves sin *ax* of various frequencies *a* = 2^*k*^, *k ∈ ℕ*. For *k* = 0, the wave crosses the *x*-axis (a zero-crossing) when *x = π*. When we double the frequency at *k* = 1, our wave is now twice as likely to cross the zero and is thus less smooth than *k* = 0. This simple zero-crossing intuition for smoothness is relatively powerful, as we will see shortly.

Next, we show that our notions of smoothness and frequency are readily applicable to data that is not regularly structured, such as single-cell data. The graph Laplacian 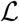 can be considered as a graph analog of the Laplace (second derivative) operator ∇^2^ from multivariate calculus. This relation can be verified by deriving the graph Laplacian from first principles.

For a graph 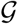 on *N* vertices, its graph Laplacian 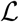 and an arbitrary graph signal f ∈ ℝ^*N*^, we use **Equation 5** to write

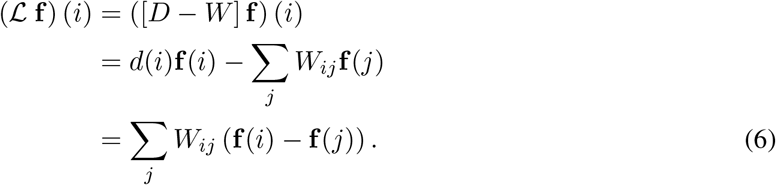

As the graph Laplacian is a weighted sum of differences of a function around a vertex, we may interpret it analogously to its continuous counterpart as the curvature of a graph signal. Another common interpretation made explicit by the derivation in **Equation 6** is that 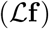(*i*) measures the *local variation* of a function at vertex *i*.

Local variation naturally leads to the notion of *total variation*,

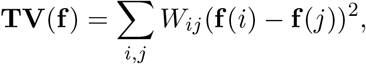

which is effectively a sum of all local variations. TV (f) describes the global smoothness of the graph signal f. In this setting, the more smooth a function is, the lower the value of the variation. This quantity is more fundamentally known as the *Laplacian quadratic form,*

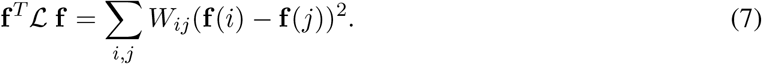

Thus, the graph Laplacian can be used as an operator and in a quadratic form to measure the smoothness of a function defined over a graph. One effective tool for analyzing such operators is to examine their eigensystems. In our case, we consider the eigendecomposition 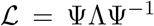, with eigenvalues^3^ Λ: = {0 = λ_1_ ≤ λ_2_ ≤ … ≤ *λ_N_*} and corresponding eigenvectors 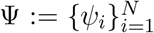. As the Laplacian is a square, symmetric matrix, the spectral theorem tells us that its eigenvectors in form an orthonormal basis for ℝ^*N*^. Furthermore, the Courant-Fischer theorem establishes that the eigenvalues in Λ are local minima of 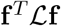 when f^*T*^f = 1 and f ∈ *U* as dim(*U*) = *i* = 1,2,.,.,*N*. At each eigenvalue λ_*i*_ this function has f = *ψ_i_*. In summary, the eigenvectors of the graph Laplacian (1) are an orthonormal basis and (2) minimize the Laplacian quadratic form for a given dimension.

Henceforth, we use the term *graph Fourier basis* interchangeably with graph Laplacian eigenvectors, as this basis can be thought of as an extension of the classical Fourier modes to irregular domains [18]. In particular, the ring graph eigenbasis is composed of sinusoidal eigenvectors, as they converge to discrete Fourier modes in one dimension. The graph Fourier basis thus allows one to define the *graph Fourier transform* (GFT) by direct analogy to the classical Fourier transform.

The GFT of a signal *f* is given by 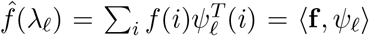, which can also be written as the matrix-vector product

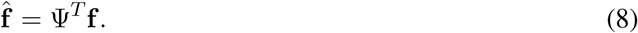

As this transformation is unitary, the inverse graph Fourier transform (IGFT) is 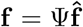. Although the graph setting presents a new set of challenges for signal processing, many classical signal processing notions such as filterbanks and wavelets have been extended to graphs using the GFT. We use the GFT to process, analyze, and cluster experimental signals from single-cell data using a novel graph filter construction and a new harmonic clustering method.

#### 4.1.5 The manifold heat filter

In the MELD algorithm, we seek to estimate the change in sample density between experimental labels along a manifold represented by a cell similarity graph. To estimate sample density along the graph, we employ a novel graph filter construction, which we explain in the following sections. To begin, we review the notion of filtering with focus on graphs and demonstrate manifold heat filter in a low-pass setting. Next, we demonstrate the expanded version of the manifold heat filter and provide an analysis of its parameters. Finally, we provide a simple solution to the manifold heat filter that allows fast computation.

#### 4.1.6 Filters on graphs

Filters can be thought of as devices that alter the spectrum of their input. Filters can be used as bases, as is the case with wavelets, and they can be used to directly manipulate signals by changing the frequency response of the filter. For example, many audio devices contain an equalizer that allows one to change the amplitude of bass and treble frequencies. Simple equalizers can be built simply by using a set of filters called a filterbank. In the MELD algorithm, we use a tunable filter to estimate density of a sample indicator signal on a single-cell graph.

Mathematically, graph filters work analogously to classical filters. Specifically, a filter takes in a signal and attenuates it according to a frequency response function. This function accepts frequencies and returns a response coefficient. This is then multiplied by the input Fourier coefficient at the corresponding frequency. The entire filter operation is thus a reweighting of the input Fourier coefficients. In low-pass filters, the function only preserves frequency components below a threshold. Conversely, high-pass filters work by removing frequencies below a threshold. Bandpass filters transfer frequency components that are within a certain range of a central frequency. The tunable filter in the MELD algorithm is capable of producing any of these responses.

As graph harmonics are defined on the set Λ, it is common to define them as functions of the form *h*: [0, max(Λ)] ↦ [0,1]. For example, a low pass filter with cutoff at λ_*k*_ would have *h*(*x*) > 0 for *x* < λ_*k*_ and *h*(*x*) = 0 otherwise. By abuse of notation, we will refer to the diagonal matrix with the filter *h* applied to each Laplacian eigenvalue as *h*(Λ), though *h* is not a set-valued or matrix-valued function. Filtering a signal f is clearest in the spectral domain, where one simply takes the multiplication 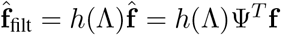.

Finally, it is worth using the above definitions to define a vertex-valued operator to perform filtering. As a graph filter is merely a reweighting of the graph Fourier basis, one can construct the *filter matrix,*

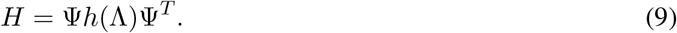

A manipulation using **Equation 8** will verify that *H*f is the WGFT of 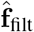. This filter matrix will be used to solve the manifold heat filter in approximate form for computational efficiency.

#### 4.1.7 Laplacian Regularization

A simple assumption for density estimation is *smoothness.* In this model the density estimate is assumed to have a low amount of neighbor to neighbor variation. *Laplacian regularization* [57–65] is a simple technique that targets signal smoothness via the optimization

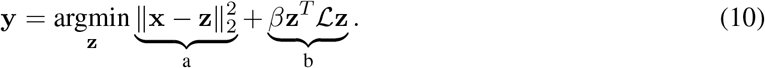

Note that this optimization has two terms. The first term (a), called a *reconstruction penalty*, aims to keep the density estimate similar to the input sample information. The second term (b) ensures smoothness of the signal. Balancing these terms adjusts the amount of smoothness performed by the filter.

Laplacian regularization is a sub-problem of the manifold heat filter that we will discuss for low-pass filtering. In the above, a reconstruction penalty (a) is considered alongside the Laplacian quadratic form (b), which is weighted by the parameter *β.* The Laplacian quadratic form may also be considered as the norm of the *graph gradient*, i.e.

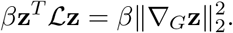

Thus one may view Laplacian regularization as a minimization of the edge-derivatives of a function while preserving a reconstruction. Because of this form, this technique has been cast as *Tikhonov regularization* [59, 66], which is a common regularization to enforce a low-pass filter to solve inverse problems in regression. In our results we demonstrate a manifold heat filter that may be reduced to Laplacian regularization using a squared Laplacian.

In **Section 4.1.6** we introduced filters as functions defined over the Laplacian eigenvalues (*h*(Λ)) or as vertex operators in **Equation 9**. Minimizing optimization **Equation 10** reveals a similar form for Laplacian regularization. Although Laplacian regularization filter is presented as an optimization, it also has a closed form solution. We derive this solution here as it is a useful building block for understanding the sample-associate density estimate. To begin,

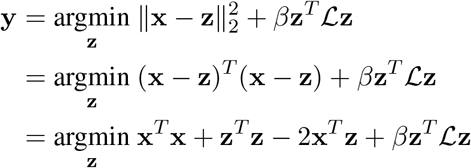

Substituting y = z, we next differentiate with respect to y and set this to 0,

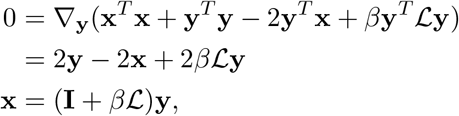

so the global minima of (10) can be expressed in closed form as

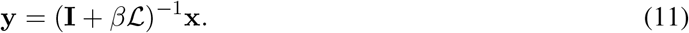

As the input x is a graph signal in the vertex domain, the least squares solution (11) is a filter matrix 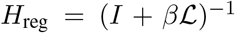 as discussed in **Section 4.1.6**. The spectral properties of Laplacian regularization immediately follow as

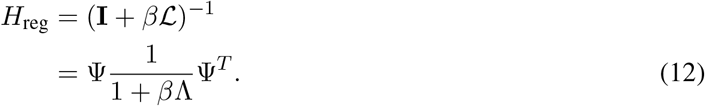

Thus Laplacian regularization is a graph filter with frequency response *h*_reg_(λ) = (1 + *β*λ)^-1^. **Figure S13b** shows that this function is a low-pass filter on the Laplacian eigenvalues with cutoff parameterized by *β*.

#### 4.1.8 Tunable Filtering

Though simple low-pass filtering with Laplacian regularization is a powerful tool for many machine learning tasks, we sought to develop a filter that is flexible and capable of filtering the signal at any frequency. To accomplish these goals, we introduce the manifold heat filter:

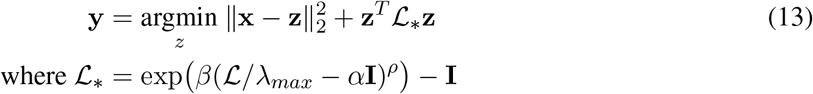

This filter expands upon Laplacian regularization by the addition of a new smoothness structure. Early and related work proposed the use of a power Laplacian smoothness matrix *S* in a similar manner as we apply here [59], but little work has since proven its utility. In our construction, *α* is referred to as modulation, *β* acts as a reconstruction penalty, and *ρ* is filter order. These parameters add a great deal of versatility to the manifold heat filter, and we demonstrate their spectral and vertex effects in **Figure S13**, as well as provide mathematical analysis of the MELD algorithm parameters in **Section 4.1.8**. Finally, in **Section 4.1.11** we discuss an implementation of the filter.

A similar derivation as **Section 4.1.7** reveals the filter matrix

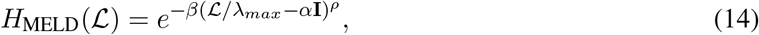

which has the frequency response

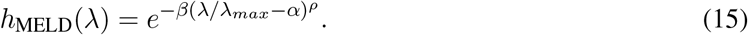

Thus, the value of the MELD algorithm parameters in the vertex optimization (**Equation 13**) has a direct effect on the graph Fourier domain.

#### 4.1.9 Parameter Analysis

*β* steepens the cutoff of the filter and shifts it more towards its central frequency (**Figure S13b**). In the case of *α* = 0, this frequency is λ_1_ = 0. This is done by scaling all frequencies by a factor of *β*. For stability reasons, we choose *β* > 0, as a negative choice of *β* yields a high frequency amplifier.

The parameters *α* and *ρ* change the filter from low pass to band pass or high pass. **Figure S13** highlights the effect on frequency response of the filters and showcases their vertex effects in simple examples. We begin our mathematical analysis with the effects of *ρ*.

*ρ* powers the Laplacian harmonics. This steepens the frequency response around the central frequency of the manifold heat filter. Higher values of *ρ* lead to sharper tails (**Figure S13c, S13e**), limiting the frequency response outside of the target band, but with increased response within the band. Finally, *ρ* can be used to make a high pass filter by setting it to negative values (**Figure S13f**).

For the integer powers, a basic vertex interpretation of *ρ* is available. Each column of 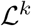 is *k-hop localized,* meaning that 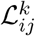 is non-zero if and only if the there exists a path length *k* between vertex *i* and vertex *j* (for a detailed discussion of this property, see Hammond et al. [54, section 5.2].) Thus, for *ρ* ∈ ℕ, the operator 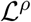 considers variation over a hop distance of *ρ*. This naturally leads to the spectral behavior we demonstrate in **Figure S13c**, as signals are required to be smooth over longer hop distances when *α* = 0 and *ρ* > 1.

The parameter *α* removes values from the diagonal of 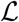. This results in a *modulation* of frequency response by translating the Laplacian harmonic that yields the minimal value for the problem (**Equation 13**). This allows one to change the central frequency, as *α* effectively modulates a band-pass filter. As graph frequencies are positive, we do not consider *α* < 0. In the vertex domain, the effect of *α* is more nuanced. We study this parameter for *α* > 0 by considering a modified Laplacian 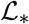 with *ρ* = 1.

To conclude, we propose a filter parameterized by reconstruction *β* (**Figure S13b**), order *ρ* (**Figure S13c, S13e**), and modulation *α* (**Figure S13d**). The parameters *α* and *β* are limited to be strictly greater than or equal to 0. When *α* = 0, *ρ* may be any integer, and it adds more low frequencies to the frequency response as it becomes more positive. On the other hand, if *ρ* is negative and *α* = 0, *ρ* controls a high pass filter. When *α* > 0, the manifold heat filter becomes a band-pass filter. In standard use cases we propose to use the parameters *α* = 0, *β* = 60, and *ρ* = 1. Other parameter values are explored further in (**Figure S14**). We note that the results are relatively robust to parameter values around this default setting. All of our biological results were obtained using this parameter set, which gives a square-integrable low-pass filter. As these parameters have direct spectral effects, their implementation in an efficient graph filter is straightforward and presented in **Section 4.1.11**.

In contrast to previous works using Laplacian filters, our parameters allow analysis of signals that are combinations of several underlying changes occurring at various frequencies. For an intuitive example, consider that the frequency of various Google searches will vary from winter to summer (low-frequency variation), Saturday to Monday (medium-frequency variation), or morning to night (high-frequency variation). In the biological context such changes could manifest as differences in cell type abundance (low-frequency variation) and cell-cycle (medium-frequency variation) [67]. We illustrate such an example in **Figure S13a** by blindly separating a medium frequency signal from a low frequency contaminating signal over simulated data. Such a technique could be used to separate low- and medium-frequency components so that they can be analyzed independently. Each of the filter parameters is explained in more detail in **Section 4.1.8**.

#### 4.1.10 Relation between MELD and the Gaussian KDE through the Heat Kernel

Kernel density estimators (KDEs) are widely used as estimating density is one of the fundamental tasks in many data applications. The density estimate is normally done in ambient space, and there are many methods to do so with a variety of advantages and disadvantages depending on the application. We instead assume that the data is sampled from some low dimensional subspace of the ambient space, e.g. that the data lies along a manifold. The MELD algorithm can be thought of as a Gaussian KDE over the discrete manifold formed by the data. This gives a density estimate at every sampled point for a number of distributions. This density estimate, as the number of samples goes to infinity, should converge to the density estimate along a continuous manifold formed by the data. The case of data uniformly sampled on the manifold was explored in [68] proving convergence of the eigenvectors and eigenvalues of the discrete Laplacian to the eigenfunctions of the continuous manifold. Coifman and Maggioni [69] explored when the data is non-uniformly sampled from the manifold and provided a kernel which can normalize out this density which results in a Laplacian modeling the underlying manifold geometry, irrespective of data density. Building on these two works, MELD allows us to estimate the manifold geometry using multiple samples with unknown distribution along it and estimate density and conditional density for each distribution on this shared manifold.

A general kernel density estimator (KDE) *f*(*x,t*) with bandwidth *t* > 0 and kernel function *K*(*x,y,t*) is defined as

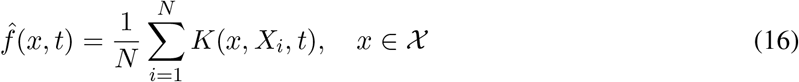

With 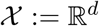, and endowed with the Gaussian kernel,

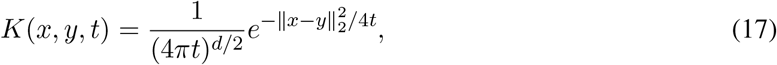

we have the Gaussian KDE in ℝ^*d*^.

This kernel is of particular interest for its thermodynamic interpretation. Namely the Gaussian KDE is a space discretization of the unique solution to the heat diffusion partial differential equation (PDE) [19, 70]:

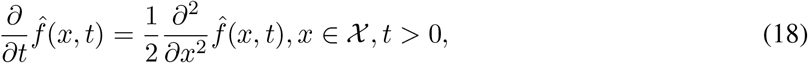

with 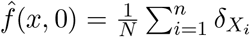 where *δ_x_* is the Dirac measure at *x*. This is sometimes called Green’s function for the diffusion equation. Intuitively, 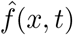 can be thought of as measuring the heat after time *t* after placing units of heat on the data points at *t* = 0.

In fact the Gaussian kernel can be represented instead in terms of the eigenfunctions of the ambient space. With eigenfunctions *φ* and eigenvalues λ, the Gaussian kernel can be alternative expressed as:

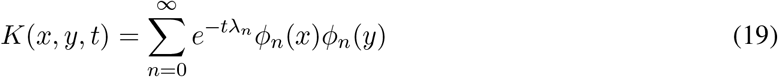

Of course for computational reasons we often prefer the closed form solution in (17). We now consider the case when 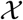 instead consists of uniform samples from a Riemannian manifold 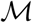 embedded in ℝ^*d*^ such that 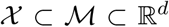. An analog to the Gaussian KDE in on a manifold is then the solution to the heat PDE restricted to the manifold, and again we can use the eigenfunction interpretation of the Green’s function in (19), except replacing the eigenfunctions of the manifold.

The eigenfunctions of the manifold can be approximated through the eigenvectors of the discrete Laplacian. The solution of the heat equation on a graph is defined in terms of the discrete Laplacian 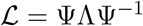 as

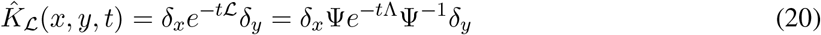

Where *δ_x_, δ_y_* are dirac functions at *x* and *y* respectively. This is equivalent to MELD when *β* = *t*λ_*max*_, *α* = 0, and *φ* = 1.

When data 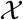 is sampled uniformly from the manifold 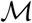 and the standard gaussian kernel is used to construct the graph, then Theorem 2.1 of Belkin and Niyogi [68] which proves the convergence of the eigenvalues of the discrete graph laplacian to the continuous laplacian implies (20) converges to the Gaussian KDE on the manifold.

However, real data is rarely uniformly sampled from a manifold. When the data is instead sampled from a smooth density 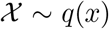 over the manifold then the density must be normalized out to recover the geometry of the manifold. This problem was first tackled in Coifman and Lafon [48], by constructing an anisotropic kernel which divides out the density at every point. This correction allows us to estimate density over the underlying *geometry of the manifold* even in the case where data is not uniformly sampled. This allows us to use samples from multiple distributions, in our case distributions over cellular states, which allows a better estimate of underlying manifold utilizing all available data.

In practice, we combine two methods to construct a discrete Laplacian that reflects the underlying data geometry over which we estimate heat propagation and perform density estimation, as explained in **Section 4.1.1**.

#### 4.1.11 Implementation

A naïve implementation of the MELD algorithm would apply the matrix inversion presented in **Equation 14**. This approach is untenable for the large single-cell graphs that the MELD algorithm is designed for, as 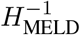 will have many elements, and, for high powers of *ρ* or non-sparse graphs, extremely dense. A second approach to solving **Equation 13** would diagonalize 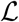 such that the filter function in **Equation 15** could be applied directly to the Fourier transform of input raw experimental signals. This approach has similar shortcomings as eigendecomposition is substantively similar to inversion. Finally, a speedier approach might be to use conjugate gradient or proximal methods. In practice, we found that these methods are not well-suited for estimating sample-associated density.

Instead of gradient methods, we use Chebyshev polynomial approximations of *h*_MELD_(λ) to rapidly approximate and apply the manifold heat filter. These approximations, proposed by Hammond et al. [54] and Shuman et al. [20], have gained traction in the graph signal processing community for their efficiency and simplicity. Briefly, a truncated and shifted Chebyshev polynomial approximation is fit to the frequency response of a graph filter. For analysis, the approximating polynomials are applied as polynomials of the Laplacian multiplied by the signal to be filtered. As Chebyshev polynomials are given by a recurrence relation, the approximation procedure reduces to a computationally efficient series of matrix-vector multiplications. For a more detailed treatment one may refer to Hammond et al. [54] where the polynomials are proposed for graph filters. For application of the manifold heat filter to a small set of input sample indicator signals, Chebyshev approximations offer the simplest and most efficient implementation of our proposed algorithm. For sufficiently large sets of samples, such as when considering hundreds of conditions, the computational cost of obtaining the Fourier basis directly may be less than repeated application of the approximation operator; in these cases, we diagonalize the Laplacian either approximately through randomized SVD or exactly using eigendecomposition, depending on user preference. Then, one simply constructs *H*_MELD_ = *h*_MELD_(Λ)ψ^*T*^ to calculate the sample-associated density estimate from the input sample indicator signals.

#### 4.1.12 Summary of the MELD algorithm

In summary, we have proposed a family of graph filters based on a generalization of Laplacian regularization framework to implement the computation of sample-associated density estimates on a graph. This optimization, which can be solved analytically, allows us to derive the relative likelihood of each sample in a dataset, as a smooth and denoised signal, while also respecting multi-resolution changes in the likelihood landscape. As we show in **Section 4.7**, this formulation performs better at deriving the true conditional likelihood in quantitative comparisons than simpler label smoothing algorithms. Further, the MELD algorithm it is efficient to compute.

The MELD algorithm is implemented in Python 3 as part of the MELD package and is built atop the `scprep, graphtools`, and `pygsp` packages. We developed `scprep` efficiently process single-cell data, and `graphtools` was developed for construction and manipulation of graphs built on data. Fourier analysis and Chebyshev approximations are implemented using functions from the `pygsp` toolbox [71].

### 4.2 Vertex-frequency clustering

Next, we will describe the vertex frequency clustering algorithm for partitioning the cellular manifold into regions of similar response to experimental perturbation. For this purpose, we use a technique proposed in Shuman et al. [21] based on a graph generalization of the classical Short Time Fourier Transform (STFT). This generalization will allow us to simultaneously localize signals in both frequency and vertex domains. The output of this transform will be a spectrogram *Q,* where the value in each entry *Q_i,j_* indicates the degree to which each sample indicator signal in the neighborhood around vertex *i* is composed of frequency *j*. We then concatenate the sample-associated relative likelihood and perform *k*-means clustering. The resultant clusters will have similar transcriptomic profiles, similar likelihood estimates, and similar *frequency trends* of the sample indicator signals. The frequency trends of the sample indicator signals are important because they allow us to infer movements in the cellular state space that occur during experimental perturbation.

We derive vertex frequency clusters in the following steps:

1. We create the cell graph in the same way as is done in **Section 4.1.1**.
2. For each vertex in the graph (corresponding to a cell in the data), we create a series of localized windowed signals by masking the sample indicator signal using a series of heat kernels centered at the vertex. Graph Fourier decomposition of these localized windows capture frequency of the sample indicator signal at different scales around each vertex.
3. The graph Fourier representation of the localized windowed signals is thresholded using a *tanh* activation function to produce pseudo-binary signals.
4. These pseudo-binarized signals are summed across windows of various scales to produce a single *N × N* spectrogram *Q*. PCA is performed on the spectrogram for dimensionality reduction.
5. The sample-associated relative likelihood is concatenated to the reduced spectrogram weighted by the L2-norm of PC1 to produce *Q* which captures both local sample indicator frequency trends and changes in conditional density around each cell in both datasets.
6. k-Means is performed on the concatenated matrix to produce vertex-frequency clusters.

#### 4.2.1 Analyzing frequency content of the sample indicator signal

Before we go into further detail about the algorithm, it may be useful to provide some intuitive explanations for why the frequency content of the sample indicator signal provides a useful basis for identifying clusters of cells affected by an experimental perturbation. Because the low frequency eigenvectors of the graph Laplacian identify smoothly varying axes of variance through a graph, we associate trends in the sample indicator signal associated these low-frequency eigenvectors as biological transitions between cell states. This may correspond to the shift in T cells from naive to activated, for example. We note that at intermediate cell transcriptomic states between the extreme states that are most enriched in either condition, we observe both low and middle frequency sample indicator signal components, see the blue cell in the cartoon in **Figure 2a**. This is because locally, the sample indicator signal varies from cell to cell, but on a large scale is varying from enriched in one condition to being enriched in the other. This is distinct from what we observe in our model when a group of cells are completely unaffected by an experimental perturbation. Here, we expect to find only high frequency variations in the sample indicator signal and no underlying transition or low-frequency component. The goal of vertex frequency clustering is to distinguish between these four cases: enriched in the experiment, enriched in the control, intermediate transitional states, and unaffected populations of cells. We also want these clusters to have variable size so that even small groups of cells that may be differentially abundant are captured in our clusters.

#### 4.2.2 Using the Windowed Graph Fourier Transform (WGFT) to identify local changes in sample indicator signal frequency

While the graph Fourier transform is useful for exploring the frequency content of a signal, it is unable to identify how the frequency content of graph signals change locally over different regions of the graph. In vertex frequency clustering, we are interested in understanding how the frequency content of the sample indicator signal changes in neighborhoods around each cell. In the time domain, the windowed Fourier transform identifies changing frequency composition of a signal over time by taking slices of the signal (e.g. a sliding window of 10 seconds) and applying a Fourier decomposition to each window independently (WFT) [56]. The result is a spectrogram *Q*, where the value in each cell *Q_i,j_* indicates the degree to which time-slice *i* is composed of frequency *j*. Recent works in GSP have generalized the constructions windowed Fourier transform to graph signals[21]. To extend the notion of a sliding window to the graph domain, Shuman et al. [21] write the operation of translation in terms of convolution as follows.

The *generalized translation operator T_i_*: ℝ^*N*^ → ℝ^*N*^ of signal *f* to vertex *i* ∈ {1,2,…,*N*} is given by

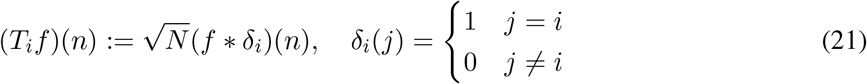

which convolves the signal *f*, in our case the sample indicator signal, with a dirac at vertex *i*. Shuman et al. [21] demonstrate that this operator inherits various properties of its classical counterpart; however, the operator is not isometric and is affected by the graph that it is built on. Furthermore, for signals that are not tightly localized in the vertex domain and on graphs that are not directly related to Fourier harmonics (e.g., the circle graph), it is not clear what graph translation implies.

In addition to translation, a *generalized modulation operator* is defined by Shuman et al. [21] as *M_k_*: ℝ^*N*^ → ℝ^*N*^ for frequencies *k* ∈ {0,1,…, *N*— 1} as

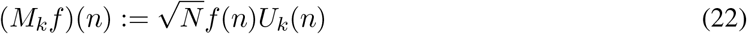

This formulation is analogous in construction to classical modulation, defined by pointwise multiplication with a pure harmonic – a Laplacian eigenvector in our case. Classical modulation translates signals in the Fourier domain; because of the discrete nature of the graph Fourier domain, this property is only weakly shared between the two operators. Instead, the generalized modulation *M_k_* translates the *DC component* of 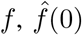, to λ_*k*_, i.e. 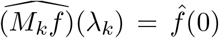. Furthermore, for any function *f* whose frequency content is localized around λ_0_, (*M_k_, f*) is localized in frequency around λ_*k*_. Shuman et al. [21] details this construction and provides bounds on spectral localization and other properties.

With these two operators, a graph windowed Fourier atom is constructed[21] for any window function *g* ∈ ℝ^*N*^

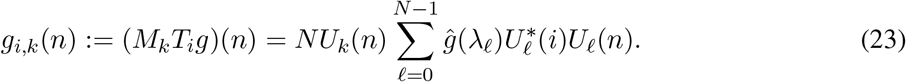

We can then build a spectrogram *Q* = (*q_ik_*) ∈ ℝ^*N* ×*N*^ by taking the inner product of each *g_i,k_*∀*i* ∈ {1, 2,…, *N*} ∧∀*k* ∈ {0,1,…, *N* — 1} with the target signal *f*

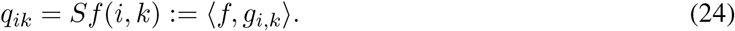

As with the classical windowed Fourier transform, one could interpret this as segmenting the signal by windows and then taking the Fourier transform of each segment

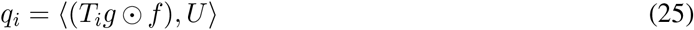

where ⊙ is the element-wise product.

#### 4.2.3 Using heat kernels of increasing scales to produce the WGFT of the sample indicator signal

To generate the spectrogram for clustering, we first need a suitable window function. We use the normalized heat kernel as proposed by Shuman et al. [21]

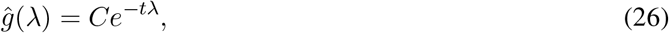

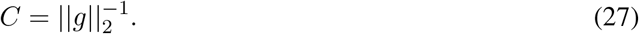

By translating this kernel, element-wise multiplying it with our target signal *f* and taking the Fourier transform of the result, we obtain a windowed graph Fourier transform of *f* that is localized based on the *diffusion distance* [21, 55] from each vertex to every other vertex in the graph.

For an input sample indicator signal f, signal-biased spectral clustering as proposed by Shuman et al. [21] proceeds as follows:

1. Generate the window matrix *P_t_*, which contains as its columns translated and normalized heat kernels at the scale *t*
2. Column-wise multiply *F_t_* = *P* ⊙ f; the *i*-th column of *F_t_* is an entry-wise product of the *i*-th window and f.
3. Take the Fourier Transform of each column of *F_t_*. This matrix, 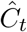 is the normalized WGFT matrix.

This produces a single WGFT for the scale *t*. At this stage, Shuman et al. [21] proposed to saturate the elements of 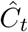 using the activation function tanh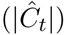 (where |.| is an element-wise absolute value). Then, k-means is performed on this saturated output to yield clusters. This operation has connections to spectral clustering as the features that k-means is run on are coefficients of graph harmonics.

We build upon this approach to add robustness, sensitivity to sign changes, and scalability. Particularly, vertex-frequency clustering builds a set of activated spectrograms at different window scales. These scales are given by simulated heat diffusion over the graph by adjusting the time-scale *t* in **Equation 26**. Then, the entire set is combined through summation.

#### 4.2.4 Combining the sample-associated relative likelihood and WGFT of the sample indicator signal

As discussed in **Section 2.4**, it is useful to consider the value of the sample likelihood in addition to the frequency content of the sample indicator. This is because if we consider two populations of cells, one of which is highly enriched in the experimental condition and another that is enriched in the control, we expect to find similar frequency content of the sample indicator signal. Namely, both should have very low-frequency content, as indicated in the cartoon in **Figure 2a**. However, we expect these two populations to have very different sample likelihood values. To allow us to distinguish between these populations, we also include the sample-associated relative likelihood in the matrix used for clustering.

We concatenate the sample-associated relative likelihood as an additional column to the multi-resolution spectrogram *Q*. However, we want to be able to tune the clustering with respect to how much the likelihood affects the result compared to the frequency information in *Q*. Therefore, inspired by spectral clustering as proposed by [72], we first perform PCA on *Q* to get *k* + 1 principle components and then normalize the likelihood by the *L*2-norm of the first principle component. We then add the likelihood as an additional column to the PCA-reduced *Q* to produce the matrix 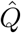. The weight of the likelihood can be modulated by a user-adjustable parameter *w*, but for all experiments in this paper, we leave *w* = 1. Finally, 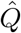 is used as input for *k*-means clustering.

The multiscale approach we have proposed has a number of benefits. Foremost, it removes the complexity of picking a window-size. Second, using the actual input signal as a feature allows the clustering to consider both frequency and sign information in the raw experimental signal. For scalability, we leverage the fact that *P_t_* is effectively a diffusion operator and thus can be built efficiently by treating it as a Markov matrix and normalizing the graph adjacency by the degree.

#### 4.2.5 Summary of the vertex frequency clustering algorithm

To identify clusters of cells that are transcriptionally similar and also affected by an experimental perturbation in the same way, we introduced an algorithm called vertex frequency clustering. Our approach builds on previous work by Shuman et al. [21] analyzing the local frequency content of the sample indicator vector as defined over the vertices of a graph. Here, we introduce two novel adaptations of the algorithm. First, we take a multiresolution approach to quantifying frequency trends in the neighborhoods around each node. By considering windowed signals that are large (i.e. contain many neighboring points) and small (i.e. very proximal on the graph), we can identify clusters both large and small that are similarly affected by an experimental perturbation. Our second contribution is the inclusion of the relative likelihood of each sample in our basis for clustering. This allows VFC to take into account the degree of enrichment of each group of cells between condition.

### 4.3 Parameter search for the MELD algorithm

To determine the optimal set of parameters for the MELD algorithm, we performed a parameter search using splatter-generated datasets. For each of the four dataset structures, we generated 10 datasets with different random seeds and 10 random ground-truth probability densities per dataset for a total of 400 datasets per combination of parameters. A coarse-grained grid search revealed that setting *α* = 0 and *ρ* =1 performed best regardless of the *β* parameter. This is expected because with these settings, the MELD filter is the standard heat kernel. A fine-grained search over parameters for *β* showed that optimal values were between 50-75 (**Figure S14**). We chose a value of 60 as the default in the MELD toolkit and this was used for all experiments. We would like to note that the optimal β parameter will vary with dataset structure and the number of cells. **Figure S14b** shows how the optimal *β* values varies as a function of the number of cells generated using splatter while keeping the underlying geometry the same.

### 4.4 Processing and analysis of the T-cell datasets

Gene expression counts matrices prepared by Datlinger et al. [13] were accessed from the NCBI GEO database accession GSE92872. 3,143 stimulated and 2,597 unstimulated T-cells were processed in a pipeline derived from the published supplementary software. First, artificial genes corresponding to gRNAs were removed from the counts matrix. Genes observed in fewer than five cells were removed. Cells with a library size higher than 35,000 UMI / cell were removed. To filter dead or dying cells, expression of all mitochondrial genes was z-scored and cells with average z-score expression greater than 1 were removed. As in the published analysis, all mitochondrial and ribosomal genes were excluded. Filtered cells and genes were library size normalized and square-root transformed. To build a cell-state graph, 100 PCA dimensions were calculated and edge weights between cells were calculated using an alpha-decay kernel as implemented in the Graphtools library (www.github.com/KrishnaswamyLab/graphtools) using default parameters. MELD was run on the cell state graph using the stimulated / unstimulated labels as input with the smoothing parameter *β* = 60. To identify a signature, the top and bottom VFC clusters by sample-associated relative likelihood were used for differential expression using a rank test as implemented in diffxpy [25] and a q-value cutoff of 0.05. GO term enrichment was performed using EnrichR using the gseapy Python package (https://pypi.org/project/gseapy/).

### 4.5 Processing and analysis of the zebrafish dataset

Gene expression counts matrices prepared by Wagner et al. [15] (the chordin dataset) were downloaded from NCBI GEO (GSE112294). 16079 cells from *chd* embryos injected with gRNAs targeting chordin and 10782 cells from *tyr* embryos injected with gRNAs targeting tyrosinase were accessed. Lowly expressed genes detected in fewer than 5 cells were removed. Cells with library sizes larger than 15,000 UMI / cell were removed. Counts were library-size normalized and square root transformed. Cluster labels included with the counts matrices were used for cell type identification.

During preliminary analysis, a group of 24 cells were identified originating exclusively from the *chd* embryos. Despite an average library size in the bottom 12% of cells, these cells exhibited 546-fold, 246-fold, and 1210-fold increased expression of Sh3Tc1, LOC101882117, and LOC101885394 respectively relative to other cells. To the best of our knowledge, the function of these genes in development is not described. These cells were annotated by Wagner et al. [15] as belonging to 7 cell types including the Tailbud – Spinal Cord and Neural – Midbrain. These cells were excluded from further analysis.

To generate a cell state graph, 100 PCA dimensions were calculated from the square root transformed filtered gene expression matrix of both datasets. Edge weights between cells on the graph were calculated using an alpha-decay kernel with parameters knn=20, decay=40. MAGIC was used to impute gene expression values using default parameters. MELD was run using the *tyr* or *chd* labels as input. The sample-associated density estimate was calculated for each of the 6 samples independently and normalized per replicate to generate 3 chordin relative likelihood estimates. The average likelihood for the chordin condition was calculated and used for downstream analysis. To identify subpopulations within the published clusters, we manually examined a PHATE embedding of each subcluster, the distribution of chordin likelihood values in each cluster, and the results of VFC subclustering with varying numbers of clusters. The decision to apply VFC was done one a per-cluster basis with the goal of identifying cell subpopulations with transcriptional similarity (as assessed by visualization) and uniform response to perturbation (as assessed by likelihood values). Cell types were annotated using sets of marker genes curated by Farrell et al. [16]. Changes in gene expression between VFC clusters was assess using a rank sum test as implemented by `diffxpy`.

### 4.6 Generation, processing and analysis of the pancreatic islet datasets

Single-cell RNA-sequencing was performed on human islet cells from three different islet donors in the presence and absence of IFN*γ*. The islets were received on three different days. Cells were cultured for 24 hours with 25ng/mL IFN*γ* (R&D Systems) in CMRL 1066 medium (Gibco) and subsequently dissociated into single cells with 0.05% Trypsin EDTA (Gibco). Cells were then stained with FluoZin-3 (Invitrogen) and TMRE (Life Technologies) and sorted using a FACS Aria II (BD). The three samples were pooled for the sequencing. Cells were immediately processed using the 10X Genomics Chromium 3’ Single-Cell RNA-sequencing kit at the Yale Center for Genome Analysis. The raw sequencing data was processed using the 10X Genomics Cell Ranger Pipeline. Raw data will be made available prior to publication.

Data from all three donors was concatenated into a single matrix for analysis. First, cells not expressing insulin, somatostatin, or glucagon were excluded from analysis using donor-specific thresholds. The data was square root transformed and reduced to 100 PCA dimensions. Next, we applied an MNN kernel to create a graph across all three donors with parameters knn=5, decay=30. This graph was then used for PHATE. MELD was run on the sample labels using default parameters. To identify coarse-grained cell types, we used previously published markers of islet cells [33]. We then used VFC to identify subpopulations of stimulated and unstimulated islet cells. To identify signature genes of IFN*γ* stimulation, we calculated differential expression between the clusters with the highest and lowest treatment likelihood values within each cell type using a rank sum test as implemented in diffxpy. A consensus signature was then obtained by taking the intersection genes with q-values < 0.05. Gene set enrichment was then calculated using gseapy.

### 4.7 Quantitative comparisons

To generate single-cell data for the quantitative comparisons, we used Splatter. Datasets were all generated using the ‘‘Paths” mode so that a latent dimension in the data could be used to create the ground truth likelihood that each cell would be observed in the ‘experimental” condition relative to the ‘control”. We focused on four data geometries: a tree with three branches, a branch and cluster with either end of the branch enriched or depleted and the cluster unaffected, a single branch with a middle section either enriched or depleted, and four clusters with random segments enriched or depleted. To create clusters, a multibranched tree was created, and all but the tips of the branches were removed. The ground truth experimental signal was created using custom Python scripts taking the “Steps” latent variable from Splatter and randomly selecting a proportion of each branch or cluster between 10% and 80% of the data was enriched or depleted by 25%. These regions were divided into thirds to create a smooth transition between the unaffected regions and the differentially abundant regions. This likelihood ratio was then centered so that, on average, half the cells would be assigned to each condition. The centered ground truth signal was used to parameterize a Bernoulli random variable and assign each cell to the experimental or control conditions. The data and sample labels were used as input to the respective algorithms.

To quantify the accuracy of MELD to approximate the ground truth likelihood ratio, we compared MELD, a kNN-smoothed signal, or a graph averaged signal to the ground truth likelihood of observing each cell in either of the two conditions. We used the Pearson’s R statistic to calculate the degree to which these estimates approximate the likelihood ratio. Each of the four data geometries was tested 30 times with different random seeds.

We also performed MELD comparisons using the T cell and zebrafish datasets described above. The preprocessed data was used to generate a three-dimensional PHATE embedding that was z-score normalized. We then used a combination of PHATE dimensions to create a ground truth probability each cell would be observed in the experimental or control condition. Cells were then assigned to either condition based on this probability as described above. We ran the same comparisons as on the simulated data with 100 random seeds per dataset.

To quantify the accuracy of VFC to detect the regions of the dataset that were enriched, depleted, or unaffected between conditions, we calculated the Adjusted Rand Score between the ground truth regions with enriched, depleted, or unchanged likelihood ratios between conditions. VFC was compared to k-Means, Spectral Clustering, Louvain, Leiden, and CellHarmony. As Leiden and Louvain do not provide a method to control the number of clusters, we implemented a binary search to identify a resolution parameter that provides the target number of clusters. Although Cell Harmony relies on an initial Louvain clustering, the tool does not implement Louvain with a tuneable resolution. It is also not possible to provide an initial clustering to CellHarmony, so we resorted to cutting Louvain at the level closest to our target number of clusters. Finally, because CellHarmony does not reconcile the disparate cluster assignments in the reference and query datasets, and because not all cells in the query dataset may be aligned to the reference we needed to generate manually new cluster labels for cells in the query dataset so that the method could be compared to other clustering tools.

To characterize the ability of MELD to characterize gene signatures of a perturbation dataset, we returned to the T cell dataset. We again used the same setup to create synthetically 3 regions with different sampling probabilities in the dataset using PHATE clusters as above. Because one of these clusters has no differential abundance between conditions, we calculated the ground truth gene expression signature between the enriched and depleted clusters only using diffxpy [25]. To calculate the gene signature for each clustering method, we performed differential expression between the most enriched cluster in the experimental condition and the most depleted cluster in the experimental condition (or highest and lowest treatment likelihood for MELD). We also considered directly performing two-sample comparison using the sample labels. To quantify the performance of each method, we used the area under the receiving operator characteristic (AUCROC) to compare the q-values produced using each method to the ground truth q-values. This process was repeated over 100 random seeds. The AUCROC curves and performance of each method relative to VFC is displayed in **Figure S6d,e**.

## 5 Data availability

Gene expression counts matrices prepared by Datlinger et al. [13] were accessed from the NCBI GEO database accession GSE92872. Gene expression counts matrices prepared by Wagner et al. [15] were downloaded from NCBI GEO accession GSE112294. The pancreatic islets datasets are available on NCBI GEO at accession GSE161465.

## 6 Code availability

Code for the MELD and VFC algorithms implemented in Python is available as part of the MELD package on GitHub https://github.com/KrishnaswamyLab/MELD and on the Python Package Index (PyPI). The GitHub repository also contains tutorials, code to reproduce the analysis of the zebrafish dataset, and code associated with several of the quantitative comparisons.

## Contributions

D.B.B., S.K., G.W., D.v.D., and A.J.G. envisioned the project. D.B.B., J.S. A.T. S.K., and G.W. developed the mathematical formulation of the problem and related numerical analysis. D.B.B, J.S., and S.G. implemented the code. D.B.B. and S.K. performed the analysis of biological and simulated data. A.L.P. and K.C.H. generated and assisted with the analysis of the pancreatic islet dataset. A.J.G. assisted with the analysis of the zebrafish data and related writing. D.B.B., J.S., A.T., S.K., and G.W. wrote the paper. S.G. assisted with the writing.

## Acknowledgements

The authors would like to thank Dr. Charles E. Vejnar, Dr. Ronald R. Coifman, Dr. James Noonan, Dr. Valerie Tornini, and Cassandra Kontur for fruitful discussions. We would like to thank Dr. Guilin Wang of the Yale Center for Genome Analysis for help in preparing the pancreatic islet data. This research was supported in part by: the Eunice Kennedy Shriver National Institute of Child Health & Human Development of the NIH (Award Number: F31HD097958) *[D.B.];* the Gruber Foundation *[S.G.];* IVADO Professor startup & operational funds, IVADO Fundamental Research Project grant PRF-2019-3583139727 [G.W.]; NIH grants R01GM135929 & R01GM130847 [G.W., S.K.]; and Chan-Zuckerberg Initiative grants 182702 & CZF2019-002440 [S.K.]. The content provided here is solely the responsibility of the authors and does not necessarily represent the official views of the funding agencies.

## 7 Supplementary Notes

### 7.1 A pipeline for analyzing single cell data using MELD

Using the MELD algorithm and VFC, it is now possible to propose a novel framework for analyzing single cell perturbation experiments. The goal of this framework is to identify populations of cells that are the most affected by an experimental perturbation and to characterize a gene signature of that perturbation. A schematic of the proposed pipeline is shown in **Figure S4**.

Prior to using the algorithms in MELD, we recommended first following established best practices for analysis of single cell data including exploratory analysis using visualization, preliminary clustering, and cluster annotation via differential expression analysis [1]. These steps ensure that the dataset is of high quality and comprises the cell types expected from the experimental setup. Following exploratory characterization, we propose the following analysis:

1. Estimate the sample-associate relative likelihood for each condition
2. Determine which exploratory clusters require subclustering with VFC by examining the likelihood distribution within each cluster, a visualization of the cluster, and the results of VFC with varying numbers of clusters
3. Create new cluster assignments using VFC
4. Annotate each cluster following best practices [1]
5. Characterize enrichment of cell populations using sample likelihood and gene signatures

The basic steps to calculate the sample-associated relative likelihood is described in **Section 2.1**. In the case of multiple replicates, we recommend calculating the sample density for each sample over a graph of all cells from all samples so long as there is sufficient overlap between samples. This overlap can be assessed using the k-nearest neighbor batch effect test described in Büttner et al. [73]. We then normalize the sample density for matched experimental and control samples of the same replicate and average across replicates to obtain an average measure of the perturbation. Variation in this likelihood across replicates can be used as a measure of consistency for the measured perturbation across cell types. The result of this step is an estimate of the probability that each cell would be observed in the treatment condition relative to the control.

Having calculated the sample-associated relative likelihood, we next recommend determining which cell populations identified during exploratory analysis require further subclustering with VFC to identify cell types enriched or depleted in the experimental condition. Determining optimal cluster resolution for single cell analysis will vary across experiments depending on the biological system being studied and the goals of each individual researcher. Instead of providing a single measure to determine the number of clusters, we outline a general strategy as a guide for users of MELD.

To determine the number of VFC clusters, we suggest taking into consideration transcriptional variation within each coarse-grained cluster and the effect of the perturbation. First, using a dimensionality reduction tool such as PHATE, examine a two or three dimensional scatter plot of the cluster colored by the sample likelihood for each cell. Here, the goal is to identify either regions that have very different likelihood values or regions of data density separated by low-density regions suggesting the present of multiple subclusters to target with VFC. We also suggest examining the distribution the likelihood values within each cluster to determine if the cells in the cluster exhibit a restricted range of responses to the perturbation or large variation that would require subclustering. Finally, we recommend running VFC with various numbers of clusters (2-5 is often sufficient) and inspecting the output on a PHATE plot and/or with a swarm plot. In ambiguous cases, it may be helpful to perform differential expression analysis and gene set enrichment to determine whether or not each cluster is biologically relevant to the experimental question under consideration [1, 74]. Importantly, not all clusters need subclustering, and we emphasize the ideal cluster resolution will vary based on the goals of each analyst.

To determine the gene signature of the perturbation, we recommend quantifying the differences in expression between VFC clusters. For experiments with only a single cell type and 3-4 VFC clusters, it is often sufficient to perform differential expression analysis between the cluster most enriched in the experimental condition and the cluster most depleted in the experimental condition. And example of this analysis is provided in **Section 2.6**. For experiments with several cell types, we recommend calculating the gene signature between the enriched and depleted VFC clusters within each exploratory cluster. To obtain a consensus gene signature, a research may take the intersection of the gene signatures within exploratory cluster. An example of this analysis is provided in **Section 2.8**.

We note that the strategy for identifying gene signatures outlined in the previous paragraph differs from the current framework employed in recent papers (**Figure S3**). Instead of comparing expression between cells from the experimental condition and the control, we compare clusters of cells identified with VFC. The rationale for the framework presented here is that if VFC clusters are transcriptionally homogeneous and exhibit a uniform response to the perturbation, we expect differences in gene expression between conditions *within* each cluster to represent biological and technical noise. However, characterizing transcriptional differences *between* cells of different clusters regardless of condition of origin will yield a description of the cell states that vary between experimental conditions. We confirm that the gene signatures obtained in this manner are more accurate than between-sample comparisons in **Section 4.7**.

### 7.2 VFC improves analysis of *chd* Cas9 knockout in zebrafish embryos

Here we provide details of our analysis of three clusters in the zebrafish datasets [15] that required further subclustering using VFC. In each example, we show biologically relevant insights that were missed in the published analysis.

The Tailbud – Presomitic Mesoderm (TPM) cluster exhibits the largest range of chordin relative likelihood values of all the clusters annotated by Wagner et al. [15]. In a PHATE visualization of the cluster, we observe many different branches of cell states, each with varying ranges of chordin relative likelihood values (**Figure 5c**). Within the TPM cluster, we find four subclusters using VFC (**Figure 5d**). Using established markers [16], we identify these clusters as immature adaxial cells, mature adaxial cells, presomitic mesoderm cells, and hematopoietic cells (**Figures 5c & S10**). Examining the distribution of chordin relative likelihood scores within each cell type, we conclude that the large range of chordin relative likelihood values within the TPM cluster is due to largely non-overlapping distributions of scores within each of these subpopulations (**Figure 5e**). The immature and mature adaxial cells, which are embryonic muscle precursors, have low chordin relative likelihood values indicating depletion of these cells in the *chd* condition which matches observed depletion of myotomal cells in chordin mutants [28]. Conversely, the presomitic mesoderm and hematopoietic mesoderm have high chordin relative likelihood values, indicating that these cells are enriched in a chordin mutant. Indeed, expansion of the hematopoietic mesoderm has been observed in chordin morphants [75] and expansion of the presomitic mesoderm was observed in siblings of the *chd* embryos by Wagner et al. [15]. This heterogeneous effect was entirely missed by the fold-change analysis, since the averaging of all cells assigned to the TPM cluster caused the depletion of adaxial cells to be masked by the expansion of the presomitic and hematopoietic mesoderm.

Another advantage of vertex-frequency clustering is that we can now differentiate between a change in gene expression levels across conditions and a change in abundance of cells expressing a given gene between conditions. When we examined marker gene expression within each of the VFC subclusters, we find different trends in expression in each cluster (**Figure 5f**). For example, Myod1, a marker of adaxial cells, is lowly expressed in the presomitic and hematopoietic mesoderm, but highly expressed in adaxial cells. Using a rank sum test, we find that Myod1 is not differentially expressed between conditions within any of the VFC clusters despite there being differential expression using all cells in the TPM cluster (**Figure 5f**). We find a similar trend with Tbx6, a mesoderm marker that is not expressed in adaxial cells. We find Tbx6 is differentially expressed between *chd* and *tyr* embryos within the whole cluster but not within the adaxial or presomitic mesoderm clusters. These results show that the observed change in expression of these genes in the published analysis was in fact due to changes in abundance of cell subpopulations that led to misleading differences in statistics calculated across multiple populations as a whole. Using the chordin relative likelihood and VFC, we can identify more appropriate clusters.

We similarly analyzed the “Epidermal – pfn1 (EPP)” and “Tailbud – Spinal Cord (TSC)” clusters which had the 6th and 3rd largest standard deviation in chordin relative likelihood values of all published clusters, respectively (**Figure S10**). We used VFC to break up the Epidermal – pfn1 cluster into two subclusters. Among the top differentially expressed genes between the resulting clusters we find tbx2b, crabp2a, and pfn1. Crabp2a, a marker of the neural plate border [16], is more lowly expressed in the cluster with higher chordin relative likelihood values, suggesting that *chd* loss-of-function inhibits expression of crabp2a. This is consistent with previous studies showing a requirement of chordin for proper gene expression patterning within the neural plate [76, 77].

Within the Tailbud – Spinal Cord cluster we further identified three subpopulations of cells using VFC. Examining gene expression within the subclusters, we can see that the published cluster contains different populations of cells. One group expresses markers of the spinal cord (neurog, elavl3) and dorsal tissues (olig3, pax6a/b) with an average chordin relative likelihood of 0.38, which is consistent with prior evidence that *chd* loss-of-function disrupts specification of the neuroectoderm and dorsal tissues such as the spinal cord [28]. Examining the two remaining subclusters, we see that these cells resemble cells found in both the TPM and Epidermal – Pfn1 clusters. One cluster exhibits high levels of crabp2a and chordin relative likelihood values <0.5 similar to the neural plate border cells subpopulation within the Epidermal – Pfn1 cluster. Similarly, we find the remaining cluster expressed markers of the tailbud and presomitic mesoderm including tbx6, sox2, and fgf8a. Together, these results demonstrate the advantage of using the sample-associated relative likelihood and vertex frequency clustering to quantify the effect of genetic loss-of-function perturbations in a complex system with many cell types.

### 7.3 Applying MELD analysis to single cell datasets with a batch effect

When jointly analyzing single cell datasets collected in different samples, difficulty may arise due to systematic changes in gene expression profiles between biologically equivalent cells [73]. These changes may be technical in nature (e.g. differences in the reverse transcription efficiency during library preparation) or biological (e.g. changes in sample preparation cause unexpected changes in biological state of otherwise equivalent cells). Regardless of the cause, the unifying feature of batch effects is that they confound the analysis a given research wants to perform. As such, it is unsurprising that dozens of batch normalization tools have been developed for single cell data [78]. However, it is important to emphasize that what constitutes a batch effect is dependent on the biological question in which a researcher is interested. Some analysts might be uninterested in variation caused by a change in cell media composition between samples, but other researchers might want to study these differences. Batch normalization tools have no way to know what variation is biologically relevant to the specific hypotheses of a given experiment and thus risk removing meaningful experimental signal when “correcting” measured values. This is problematic for analysis using MELD, because the goal of the toolkit is to quantify the differences that exist between samples without regard for the specific interests of given hypothesis. As such, we do not recommend using batch correction along the experimental axis (i.e. between experimental and control conditions) before running MELD. However, recognizing that in some cases batch correction is essential, we describe several considerations for performing MELD analysis on batch-corrected data.

For the MELD algorithm to accurately estimate relative likelihood for each sample, we assume that the graph learned from single cell data approximates the underlying cell state manifold. In **Section 20** we describe the use of an anisotropic kernel that normalizes for varying sampling density across cell states. However, some batch correction methods, such as mutual nearest neighbors [47], rely on the construction of a graph with artificially inflated weights between nodes from different samples. This graph no longer models the cell states an experiment measured, but rather enforces similarities between cells based on the heuristic of the chosen normalization model. We provide no theoretical guarantees that a graph learned from batch corrected data will accurately model the underlying probability densities of each condition.

In practice when analyzing islet cells collected from multiple donors, that applying batch correction methods across the donor label improves our ability to capture a signal of IFNg stimulation (**Section 2.8**). It is important to note that in this case, batch correction applied to a label that is orthogonal to the experimental axis. We have no examined the accuracy of the MELD algorithm when batch correction is applied between experimental and control samples, although it is our expectation that this will likely remove biological signal. We recommend any user considering applying batch correction methods prior to running MELD analysis follow these steps:

1. To determine if a batch effect exists, confirm that cells from one sample are not finding appropriate neighbors in another following the strategy outlined by Büttner et al. [73].
2. To characterize the effect, identify which genes change the most between the samples
3. Confirm that the genes that are different are not relevant to the biological question under investigation
4. Apply batch correction
5. Confirm that relevant biological differences are still present using MELD analysis
6. If the biological differences are not present, repeat from step 1 with less batch correction. If you hit your personal recursion limit, consider that you don’t actually want to do batch correction
7. If biological differences are present, then confirm that previous batch effect has been corrected and proceed to downstream analysis

**Table S1:** Quantitative comparison of methods for label smoothing over a graph. 40 random seeds were used for each of 4 synthetic datasets. 100 random seeds were used to create sample assignments on the T cell and zebrafish datasets. Average Pearson Correlation with ground truth signal is displayed with standard deviation in parentheses. Top performing algorithm is bolded.

| Dataset | Rel. Likelihood | Graph Averaging | kNN Averaging |
| --- | --- | --- | --- |
| Branch and Cluster | **0.82 (0.05)** | 0.41 (0.05) | 0.73 (0.04) |
| Non-monotonic | **0.94 (0.03)** | 0.52 (0.06) | 0.85 (0.03) |
| Four clusters | **0.91 (0.06)** | 0.44 (0.07) | 0.76 (0.07) |
| Three Branches | **0.90 (0.03)** | 0.48 (0.07) | 0.73 (0.07) |
| T cells [13] | **0.98 (0.01)** | 0.72 (0.06) | 0.32 (0.04) |
| Zebrafish [15] | **0.98 (0.01)** | 0.53 (0.07) | 0.80 (0.07) |

**Table S2:** Quantitative comparison of clustering methods to identify the cell types affected by a simulated experimental perturbation using real world data.

| Dataset | VFC | Spectral | Louvain | Leiden | KMeans | CellHarmony |
| --- | --- | --- | --- | --- | --- | --- |
| T cell [13] | **0.62 (0.07)** | 0.23 (0.11) | 0.31 (0.13) | 0.34 (0.14) | 0.11 (0.04) | 0.13 (0.05) |
| Zebrafish [15] | **0.53 (0.31)** | 0.13 (0.15) | 0.23 (0.22) | 0.19 (0.21) | 0.23 (0.20) | 0.22 (0.16) |

**Figure S1:**
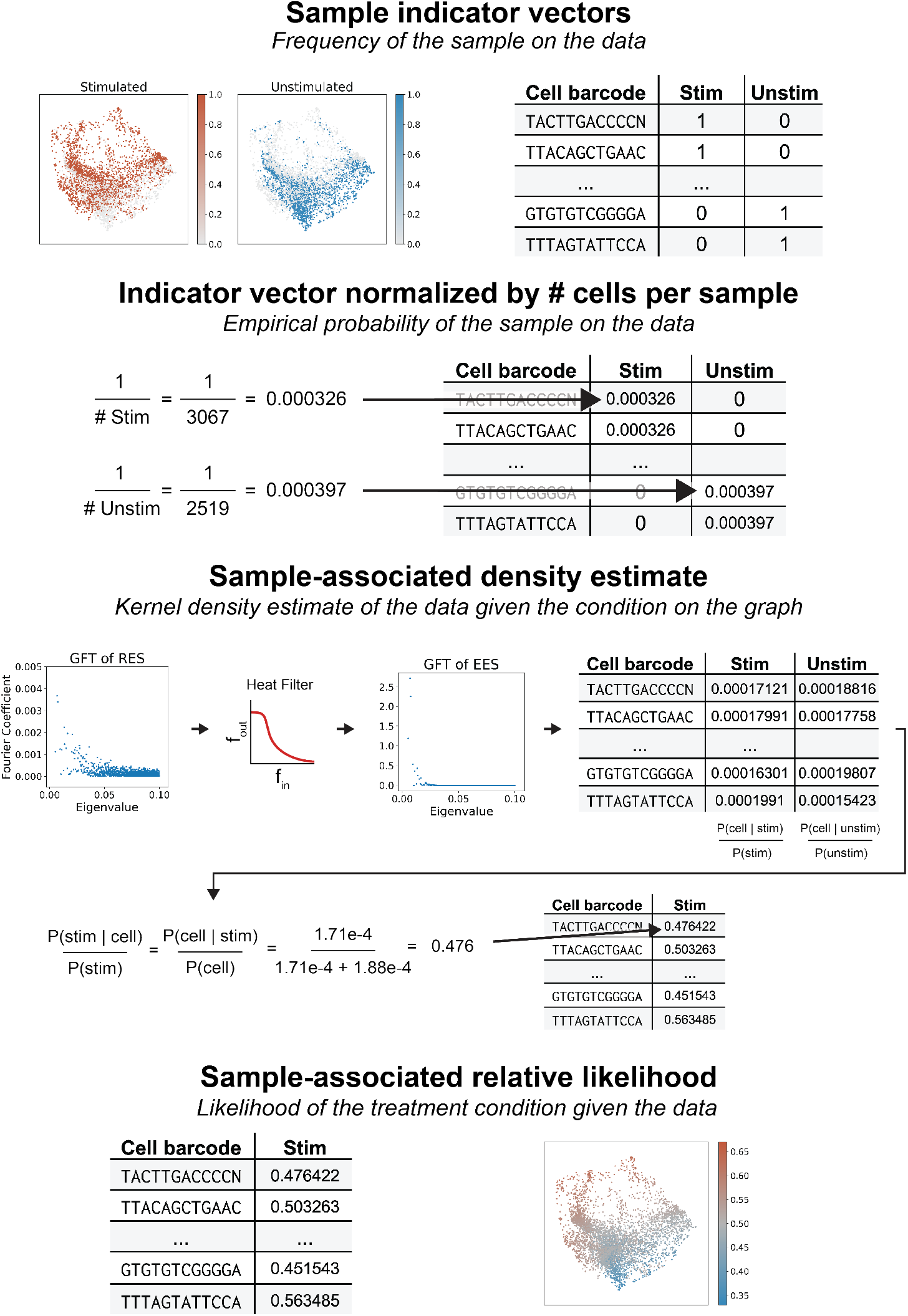
A step-by-step visual representation of the sample-associated relative likelihood algorithm using data from Datlinger et al. [13]. The sample labels are used to create a one-hot indicator signal for each condition. These one-hot signals are then column-wise L1-normalized such that the sum of each vector is 1. This gives each sample equal weight over the manifold despite a potential uneven number of cells in each condition. Next, the manifold heat filter is used to calculate a kernel density estimate for each condition. These sample-associated density estimates are then row-wise L1-normalized to yield the relative likelihood that each cell would be observed in each condition. The relative likelihood of the treatment condition relative to the control is used for two-condition experiments.

**Figure S2:**
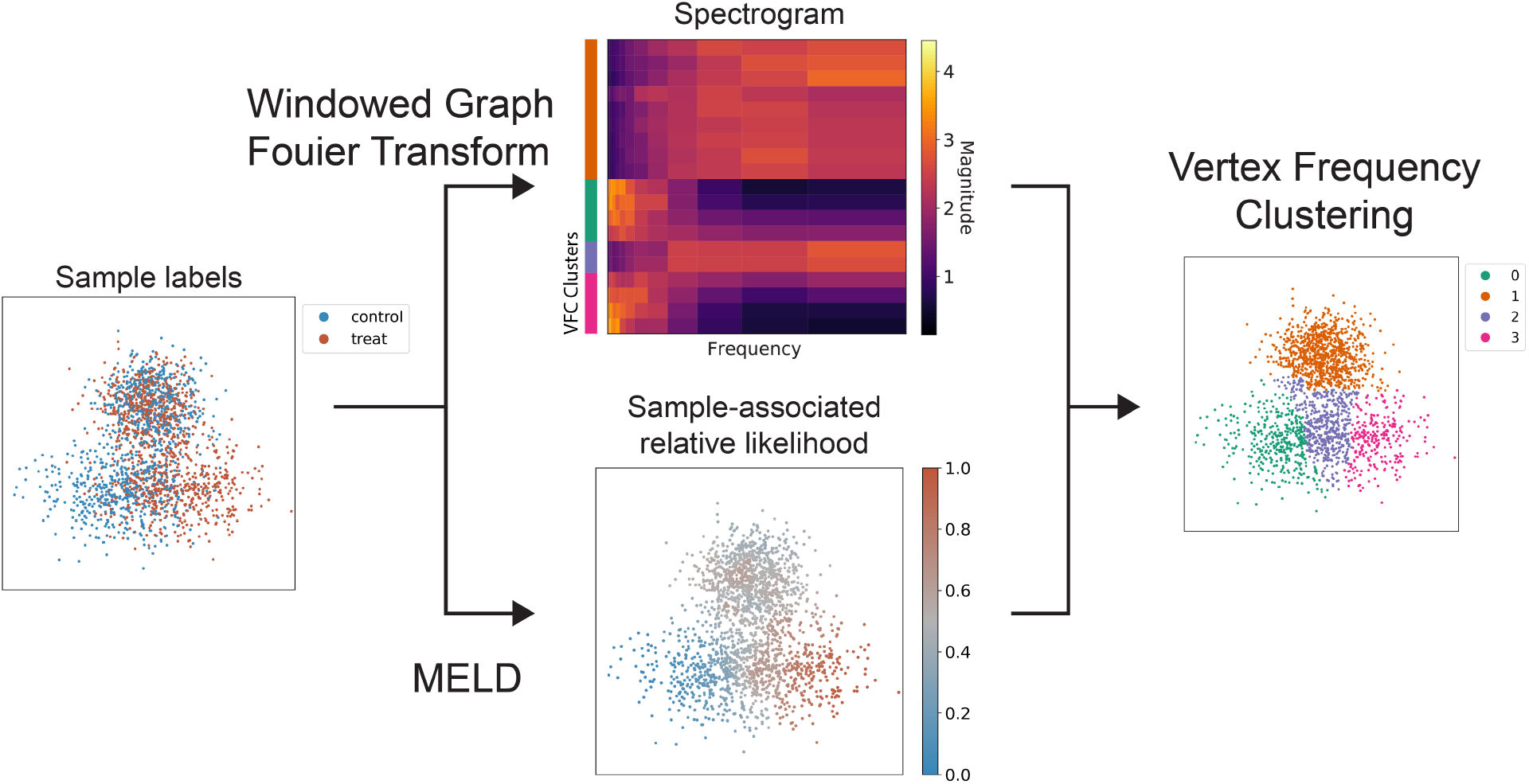
Vertex-Frequency clustering with MELD. A Gaussian mixture model was used to generate N = 2000 points in a mixture of three Gaussian distributions. This experiment is representative of a two-cell type experiment (split by Dim 2) in which one sample changes (bottom clusters) along Dim 1 due to the experiment while the other remains mixed (top clusters). Briefly, the sample labels (left) are used for (1) a windowed graph Fourier Transform to obtain vertex-frequency information (above, logarithmically downsampled for clarity) and (2) to calculate the sample-associated relative likelihood. These measures are concatenated together and clustered with *k*-Means. The clusters (right) separate the two groups of data (orange and green/purple/pink), and finds a separate grouping of points that are in transition from green to pink, shown in purple. One may see along the left side of the spectrogram that points in the green and pink clusters are found on relatively low frequency patterns with high activations in lower frequencies, whereas the transition group in purple has a well-separated medium frequency pattern. The well-mixed, nonresponsive population is entirely high frequency.

**Figure S3:**
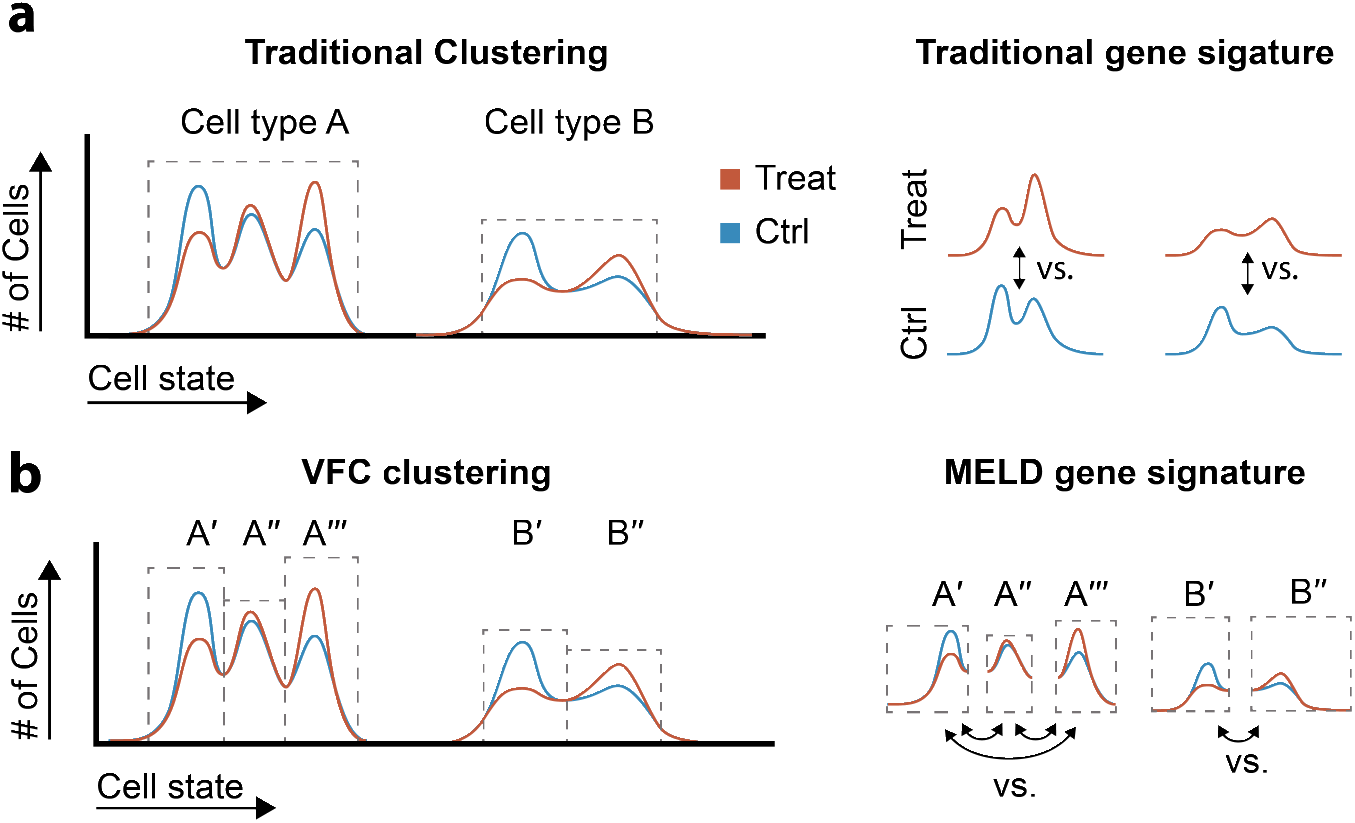
Identifying gene signatures using MELD. (**a**) In traditional gene signature analysis, clusters are identified based on data geometry and may not capture subpopulations of cells with varying response to a perturbation. In this framework, gene signatures are calculated by comparing cells from the experimental and control condition within each cluster. (**b**) To identify gene signatures of a perturbation with the MELD toolkit, we propose first partitioning cell populations with divergent responses to an experimental perturbation prior to differential expression analysis. We then assume that the differences within each VFC cluster is noise. Differential expression can either be calculated between subclusters identified by VFC (as shown) or by comparing each VFC cluster to the rest of the dataset independently.

**Figure S4:**
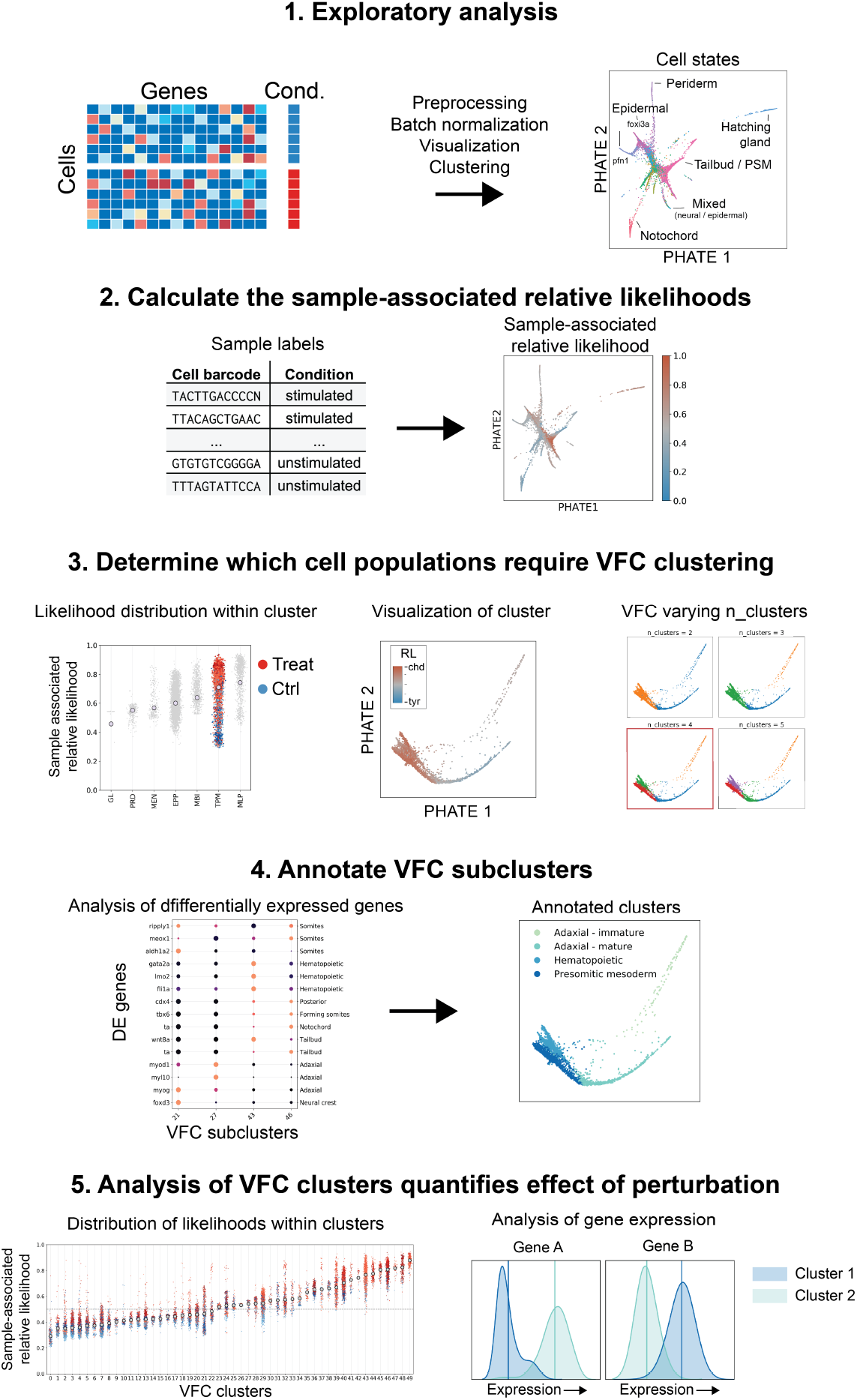
Overview of a pipeline for single cell analysis using MELD. (**1.**) Initial exploratory analysis of the dataset should follow established best practices to identify coarse-grained cell populations [1, 74]. (**2.**) Calculating the sample-associated relative likelihood provides a measure for each cell describing the probability that cell would be observed in the experimental condition relative to the control. (**3.**) To identify populations most affected by a perturbation, we consider several sources of information regarding biological heterogeneity and the effect of the perturbation within each exploratory cluster. We then apply VFC at the determined cluster resolution. (**4.**) To assess the biological relevance of each VFC cluster, standard methods for cluster annotation can be applied. (**5.**) To characterize the gene signature of the perturbation, we compare expression differences between VFC clusters with varying relative likelihood distributions.

**Figure S5:**
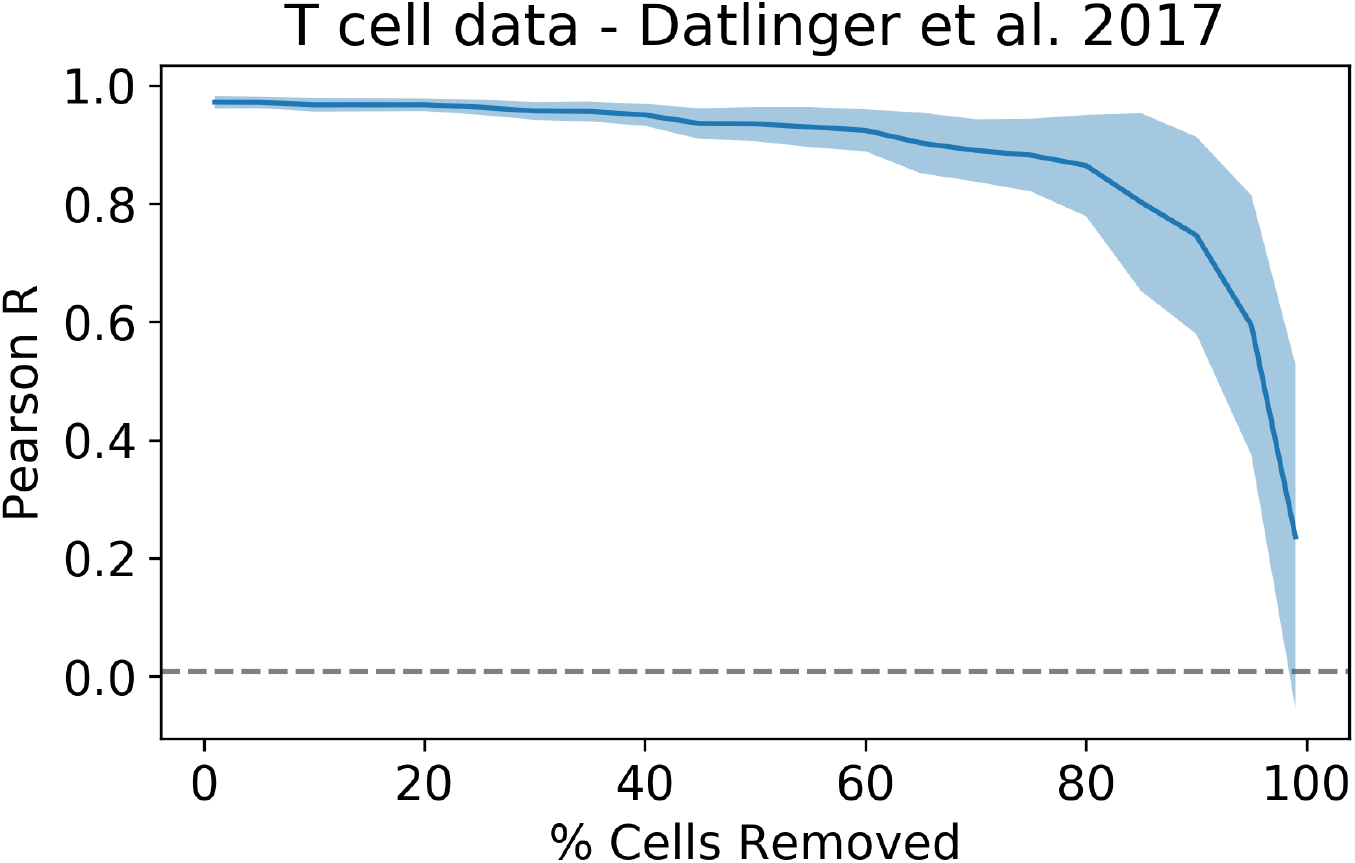
Result of down-sampling on accurately recovering simulated relative likelihood values. Using the procedure described in **Section 4.7**, we generated 100 random ground truth relative likelihoods and then removed between 1-99% of the cells in the dataset before running the MELD with default parameters. The average Pearson’s R is shown as a function of the number of cells removed prior to estimating the sample-associated relative likelihood. The shaded area demarks ±1 standard deviation. We observe an average correlation >0.9 for all experiments with at least 35% of the data present, or 1956 out of 5591 cells.

**Figure S6:**
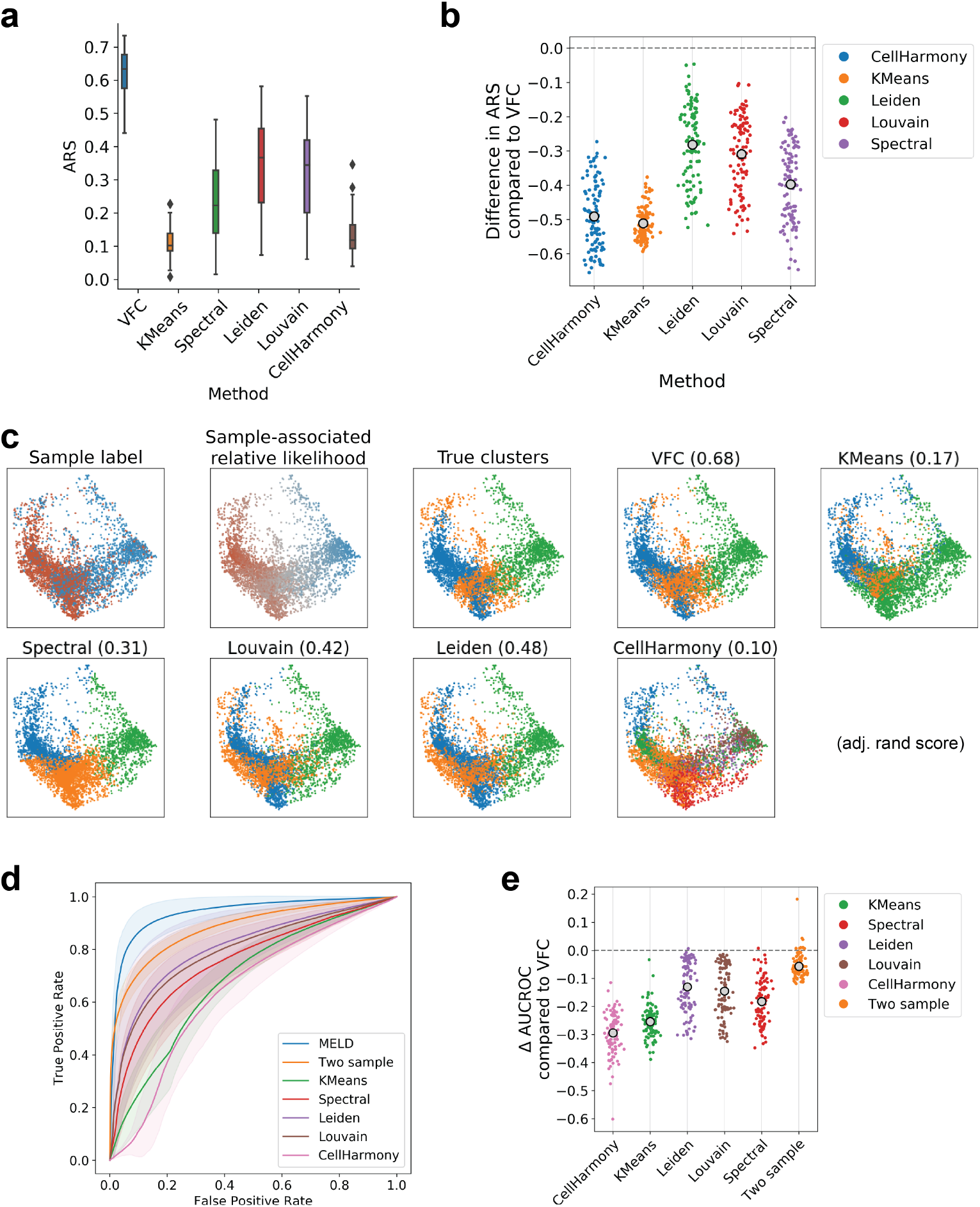
VFC accurately identifies cell populations affected by a perturbation in T cell data from Datlinger et al. [13]. (**a**) To create ground truth clusters, we artificially enriched and depleted various cell populations in either the experimental or control condition. Here we show the Adjusted Rand Score (ARS) over 100 simulations for 6 methods. For ARS, values close to 1 indicate perfect correspondence with ground truth, and values close to 0 indicate random labelling. VFC is the top performing method. (**b**) Because each simulation produced varying ARS scores for each method due to random seeds, we also consider the difference on performance between each method and VFC on each simulation. In none of 100 random seeds did any method outperform VFC. (**c**) The sample labels, sample-associated relative likelihoods, and clustering results for one randomly selected simulation. (**d**) Receiver operating characteristic (ROC) curves for the gene expression signatures described in Section 4.7. The Area Under the Curve of the ROC (AUCROC) indicates the overall performance of each strategy for identifying a gene signature. MELD is the top performing approach followed by direct comparison of the two samples. (**e**) As above, we consider the difference in AUCROC over each of 100 simulations between MELD and each method. In only 4 simulations does another method outperform MELD by more than 0.01.

**Figure S7:**
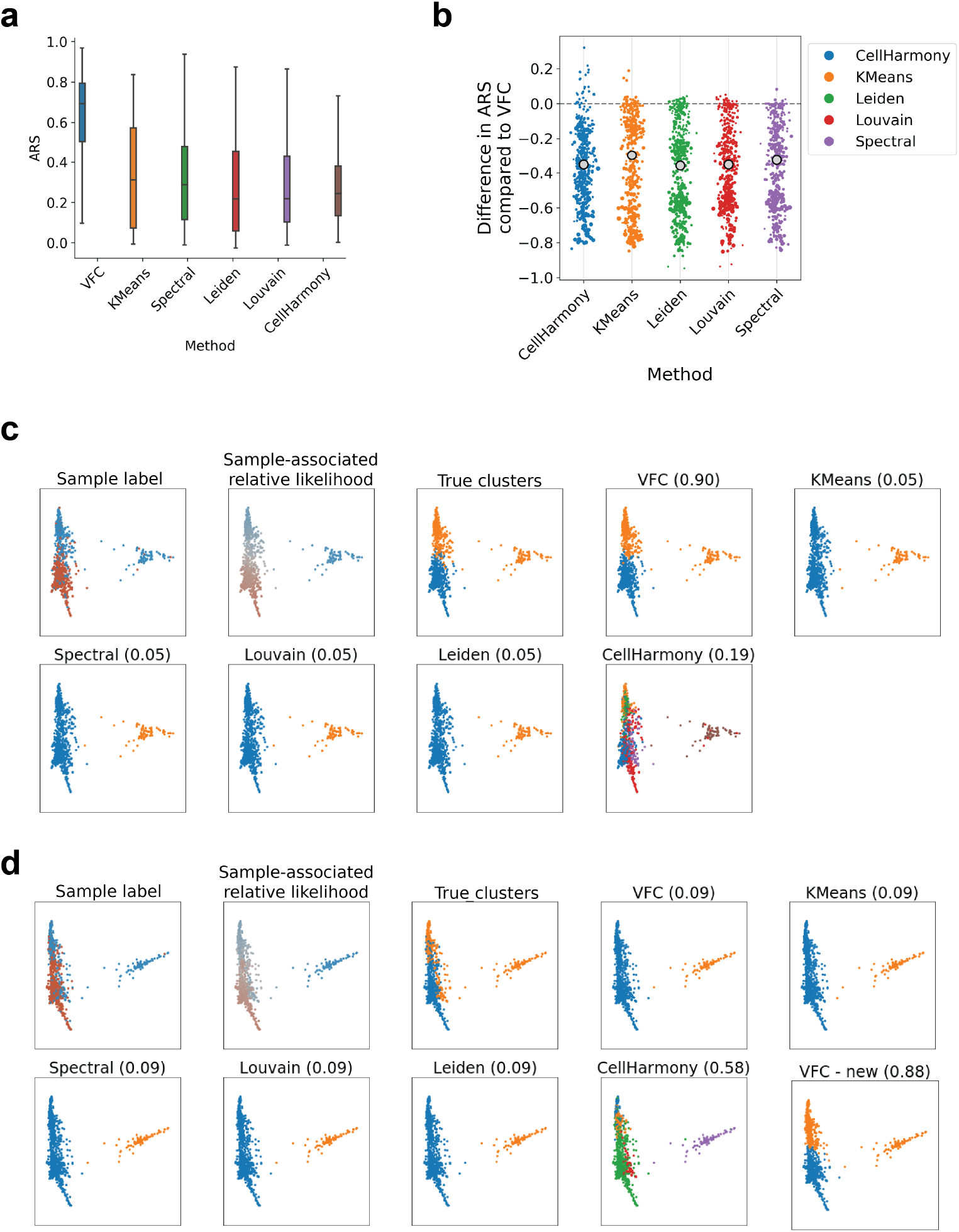
Quantitative comparison of clustering algorithms using zebrafish data from Wagner et al. [15]. (**a**) To create ground truth clusters, we artificially enriched and depleted various cell populations in either the experimental or control condition. Here we show the Adjusted Rand Score (ARS) over 100 simulations for 6 methods. VFC is the top performing method on average. (**b**) Difference on performance between each method and VFC on each simulation. (**c**) The sample labels, sample-associated relative likelihoods, and clustering results for the simulation in which VFC performed best relative to other methods and (**d**) for the simulation in which VFC performed worst relative to other methods. We found that by adjusting the weighting of the sample-associated relative likelihood from 1 (default) to 2, VFC becomes the top performing algorithm on this case (‘VFC – new’).

**Figure S8:**
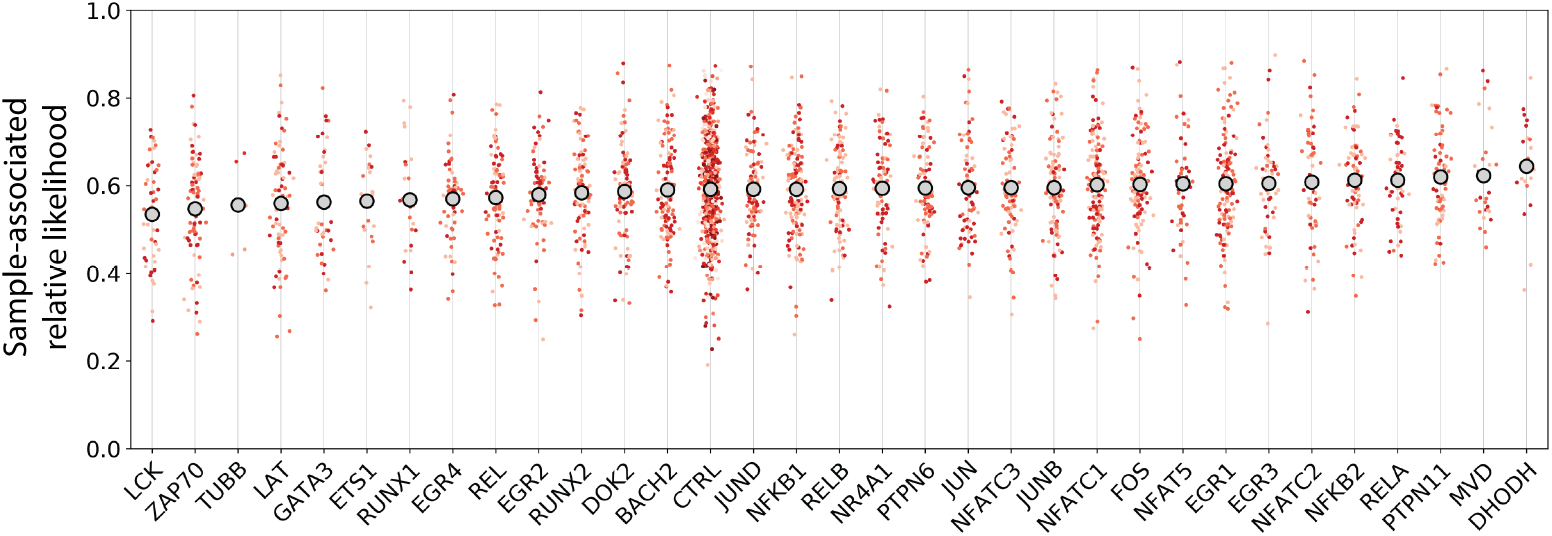
Quantitative analysis of Cas9 perturbations in T cells [13] using the MELD. Each plot shows the distribution of sampleassociate relative likelihood values for all stimulated cells transfected with gRNAs targeting a specific gene. The shade of each cell indicates the different gRNAs targeting the same gene. To determine the impact of the gRNA on the TCR activation pathway, we rank each gene by the average stimulation likelihood value. We observed a large variation in the impact of each gene knockout consistent with the published results from Datlinger et al. [13]. Encouragingly, our results agree with their bulk RNA-seq validation experiment showing greatest depletion of TCR response with knockout of kinases LCK and ZAP70 and adaptor protein LAT. We also find a slight increase in stimulation likelihood values (and therefore stimulation) in cells in which negative regulators of TCR activation are knocked out, including PTPN6, PTPN11, and EGR3.

**Figure S9:**
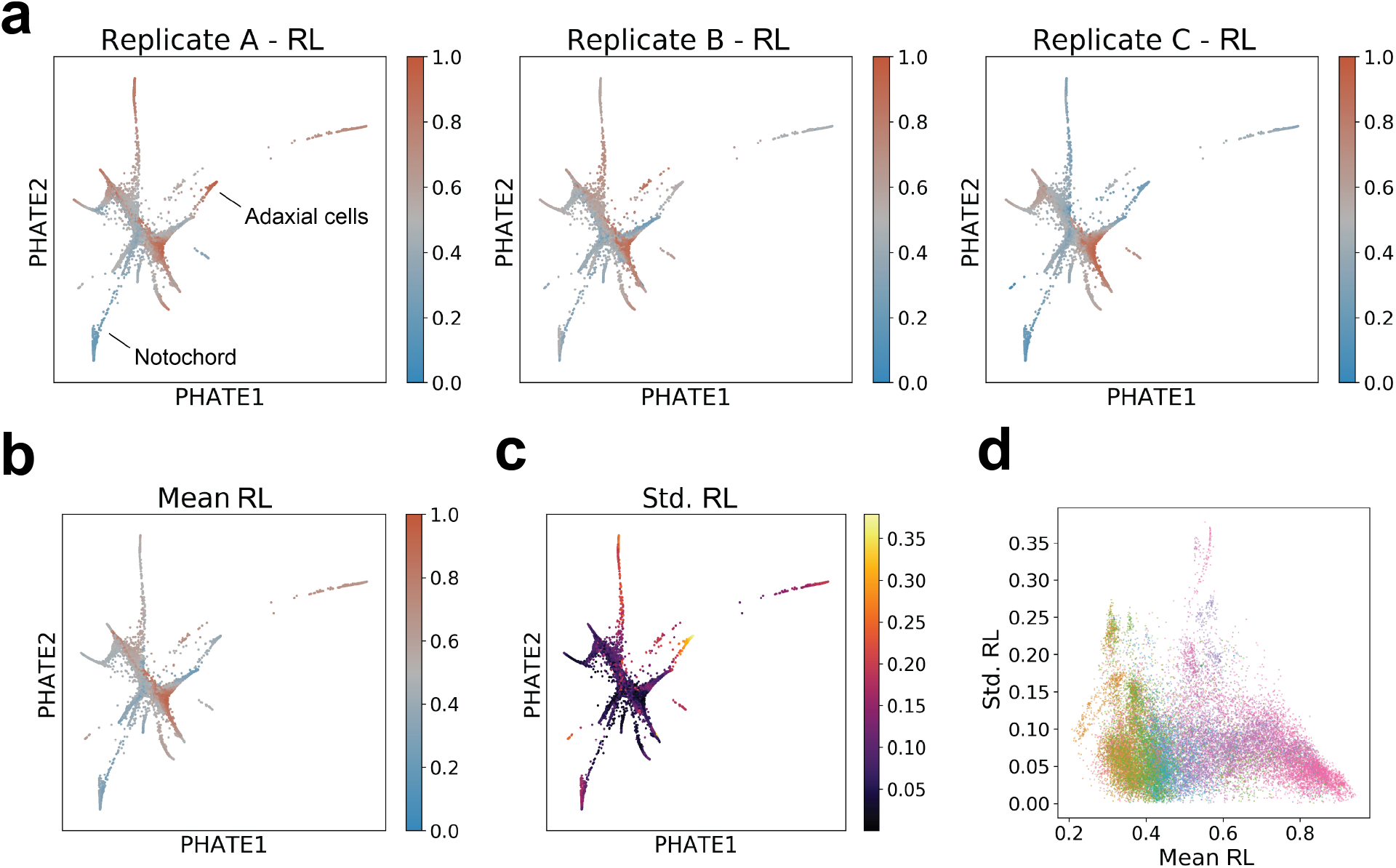
Analysis of replicates within the zebrafish data generated by Wagner et al. [15]. (**a**) Because the sample-associated relative likelihood (RL) is calculated by independently filtering a one-hot indicator vector for each condition, to calculate the chordin likelihood for each replicate, we simply row-normalize the smoothed vectors for the two signals indicating matched experimental / control pairs. For example, the ‘‘Replicate A – RL” is calculated by normalizing the “chdA” and “tyrA” filtered indicator vectors. We notice comparing replicates that the chordin likelihood for a given cell population may vary. For example, the Adaxial cell population in enriched in the Chd condition in Replicate A, but depleted in Replicate C. Similarly, cells in the Notochord population are depleted in the Chd condition in Replicates A and C, but show minimal change in abundance in Replicate B. (**b**) The average relative likelihood across all replicates is shown for each cell on a PHATE embedding. (**c**) The standard deviation of the sample-associated relative likelihood across all replicates is shown for each cell on a PHATE embedding. Regions that have higher values exhibit greater variation in their response to the experimental perturbation. We should trust the average relative likelihood values for these cells less than for cells with little variation in relative likelihood values. (**d**) A biaxial scatter plot showing the relationship between mean and standard deviation in the relative likelihood for each cell. Color indicates the cluster labels from **Figure 5a** We observe that for cells with the highest relative likelihood, the standard deviation is smaller than for cells with relative likelihood values close to 0.5 creating a slight negative Pearson correlation of −0.18.

**Figure S10:**
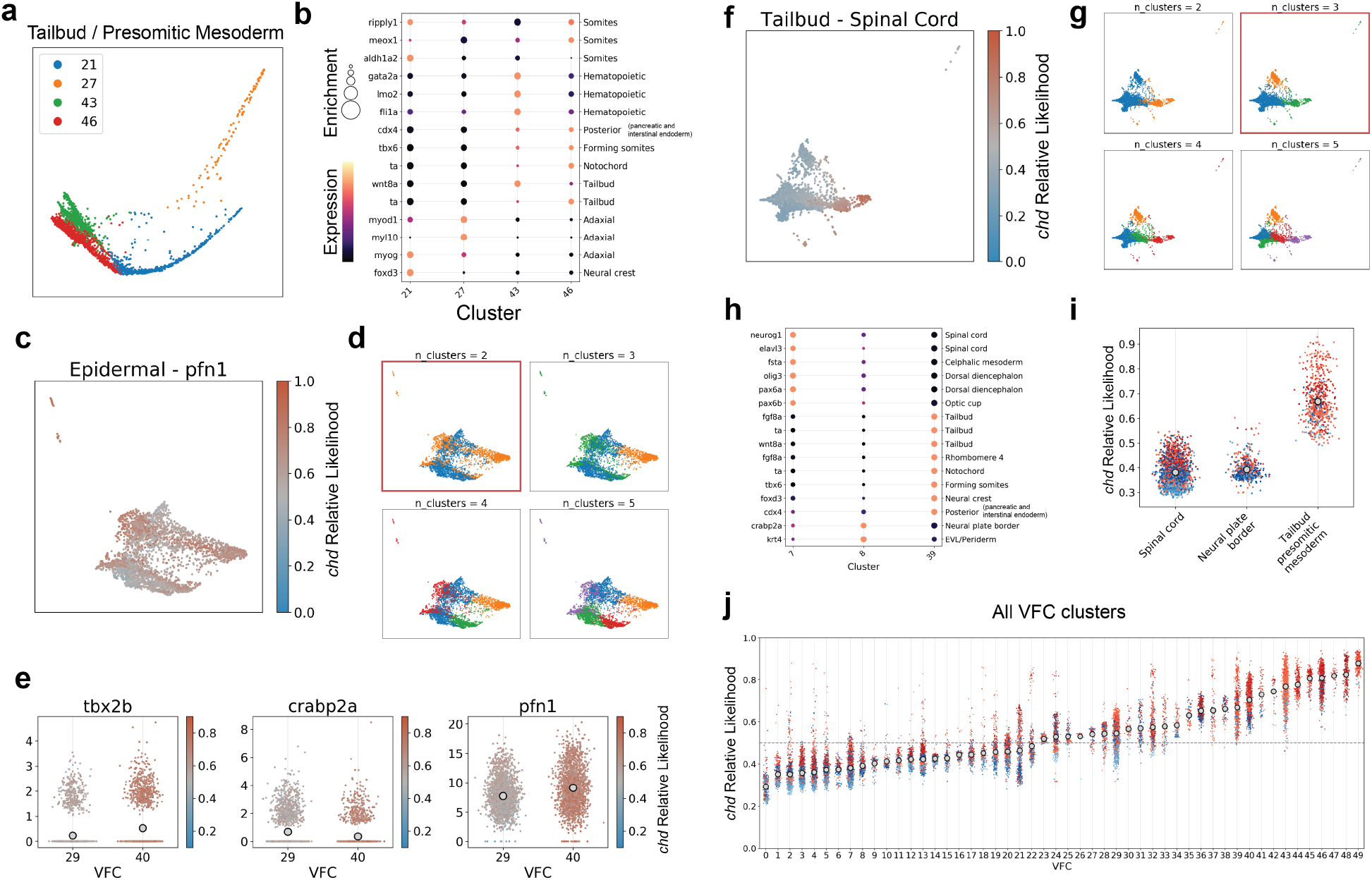
Characterization of vertex-frequency clusters in the zebrafish dataset. (**a**) Raw vertex-frequency cluster assignments on a PHATE visualization of the Tailbud – Presomitic Mesoderm cluster. (**b**) Normalized expression of previously identified marker genes of possible subtypes of the Tailbud – Presomitic Mesoderm [16]. The color of the dot for each gene in each cluster indicates the expression level and the size of the dot corresponds to the normalized Wasserstein distance between expression within cluster to all other clusters. (**c**) Distribution of chordin relative likelihood values within the ‘‘Epidermal – pfn1” cluster identified by Wagner et al. [15] shown on a PHATE plot. (**d**) Four different values of “n_clusters” that was used to create different VFC clusters with the “Epidermal – pfn1” cluster. We selected n_clusters = 2 because this identified a population of cells with similar chordin relative likelihood values and localization on the PHATE embedding. (**e**) Expression of three significantly differentially expressed genes between the two VFC subpopulations detected in the “Epidermal – pfn1” population. Tbx2b and Crabp2a were identified as markers of the epidermis and neural plate border respectively by Farrell et al. [16]. Because we observed differential expression of these two markers between the VFC subclusters suggests the ‘Epidermal – pfn1” cells identified by Wagner et al. [15] actually comprises cells originating from two distinct cell populations. (**f**) Distribution of chordin relative likelihood values within the “Tailbud – Spinal Cord” cluster identified by Wagner et al. [15] shown on a PHATE plot. (**g**) Four different values of n_clusters that was used to create different VFC clusters within the “Tailbud – Spinal Cord” cluster. We selected n_clusters = 3 because this identified populations of cells with similar likelihood values and localization on the PHATE embedding. (**h**) Same plot as in (**b**) for the subclusters of the “Tailbud – Spinal Cord”. (**i**) Distribution of relative likelihood values within each VFC subcluster show that the three subclusters are biologically distinct with differing responses to the experimental perturbation. (**j**) Repeating the VFC subclustering process for all cells, we identified a total of 50 clusters within the zebrafish dataset generated by Wagner et al. [15]. Compared to the plot in **Figure 5b**, we observed a more restricted distribution of chordin relative likelihood values within each cluster suggesting these labels represent populations of cells that are more homogeneous with respect to the experimental perturbation.

**Figure S11:**
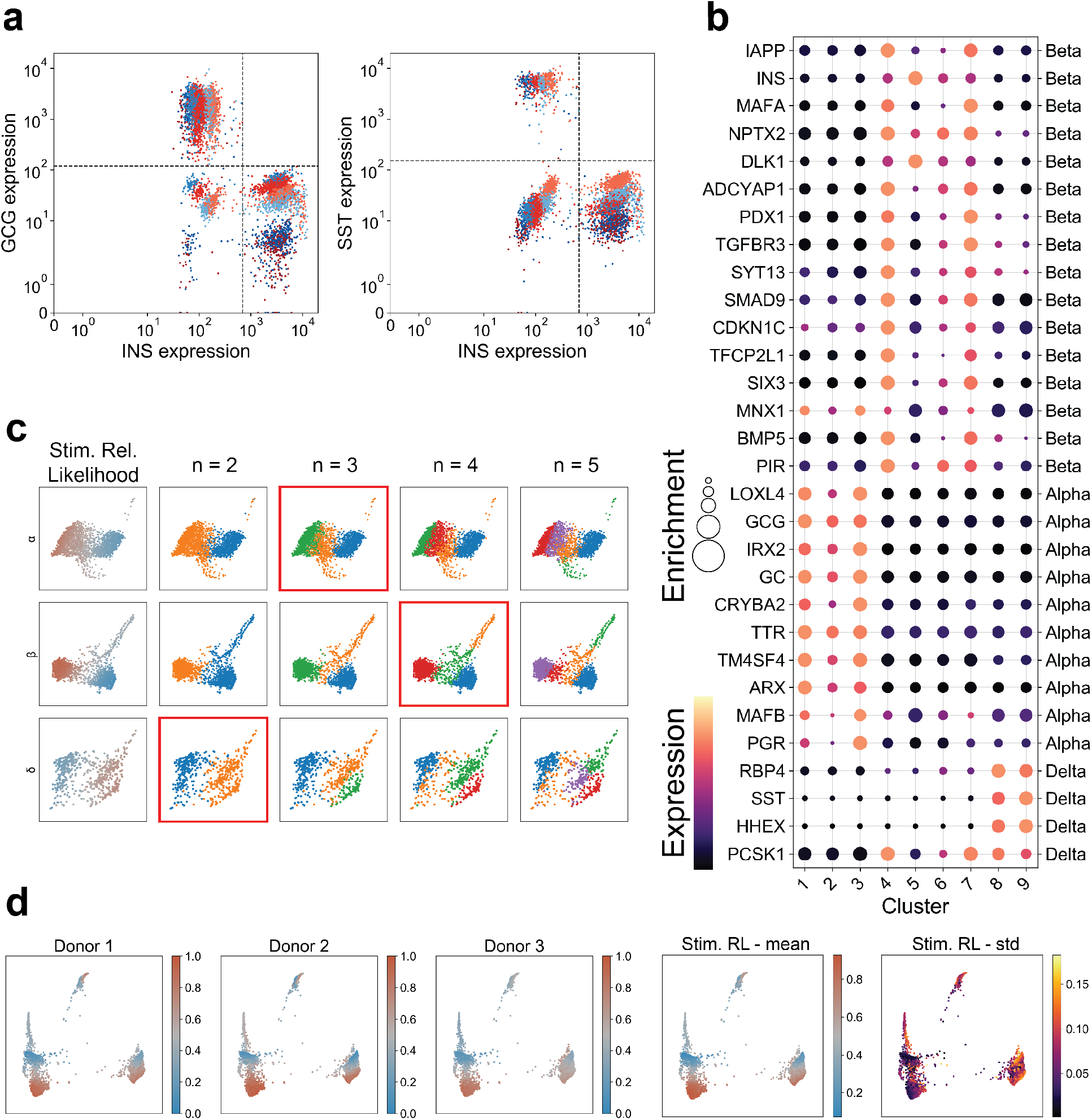
Analysis of pancreatic islet cells from three donors. (**a**) Library-size normalized expression of insulin (INS), glucagon (GCG), and somatostatin (SST) shows donor-specific batch effect across islet cells. (**b**) Normalized expression of previously identified marker genes of alpha, beta, and delta cells[33] in each cluster. The color of the dot for each gene in each cluster indicates the expression level after MAGIC and the size of the dot corresponds to the normalized Wasserstein distance between expression within cluster to all other clusters. (**c**) Results of VFC using varying numbers of clusters for each of the three cell types. The red box denotes the selected level of clustering for each cell type. (**d**) The sample-associated relative likelihood is calculated independently for each donor and then averaged to obtain the stimulated relative likelihood used in the main analysis. We also calculate the standard deviation of the relative likelihood for each cell.

**Figure S12:**
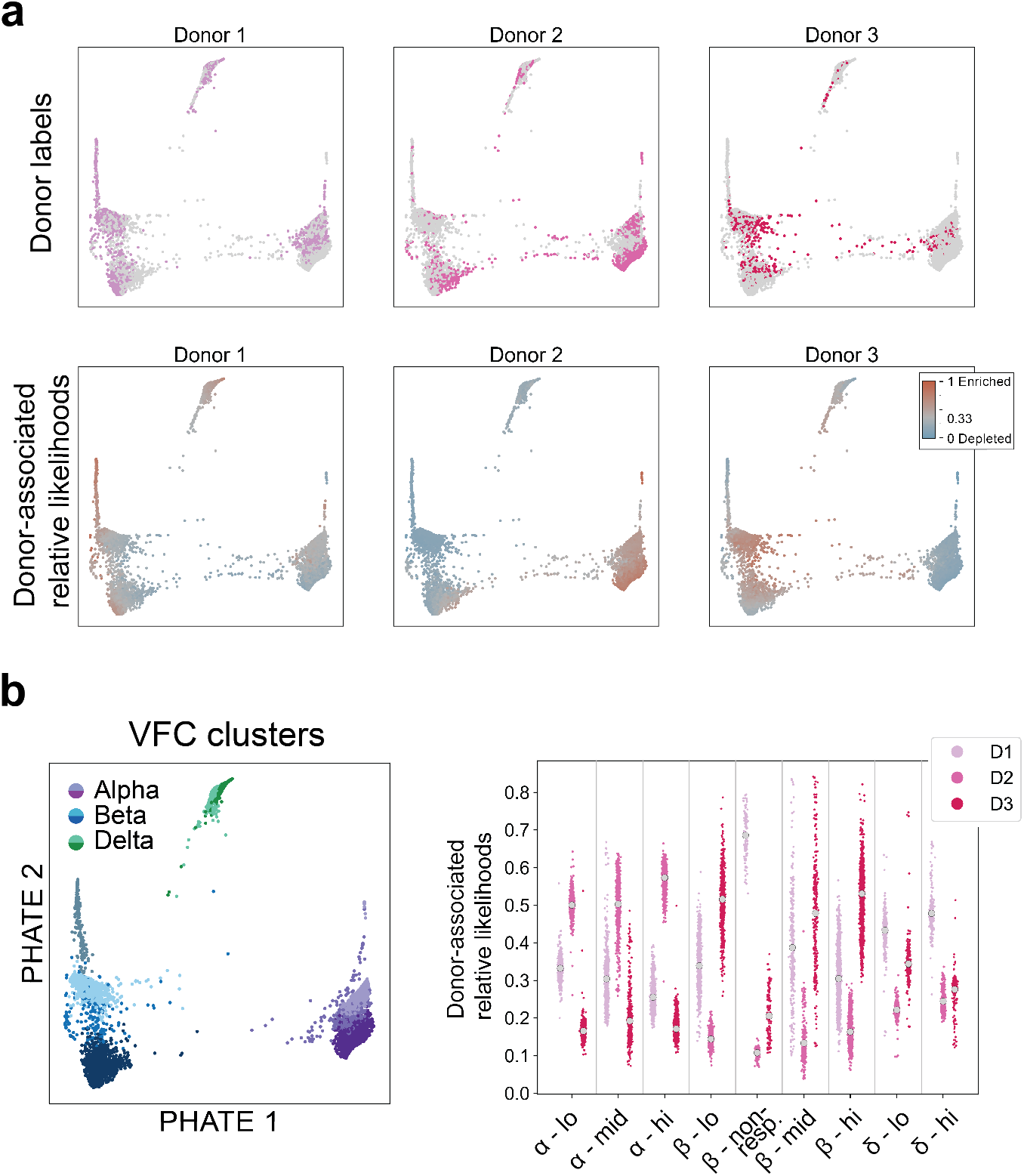
Analysis of islet cell profiles across donors. (**a**) The sample labels and sample-associated relative likelihood associated with each donor from which islet cells were obtained. (**b**) Comparison of the donor likelihood values within each vertex frequency cluster identifies changes in enrichment for each cluster in various donors. For example, the *β* – non-responsive cluster is strongly enriched in donor 1.

**Figure S13:**
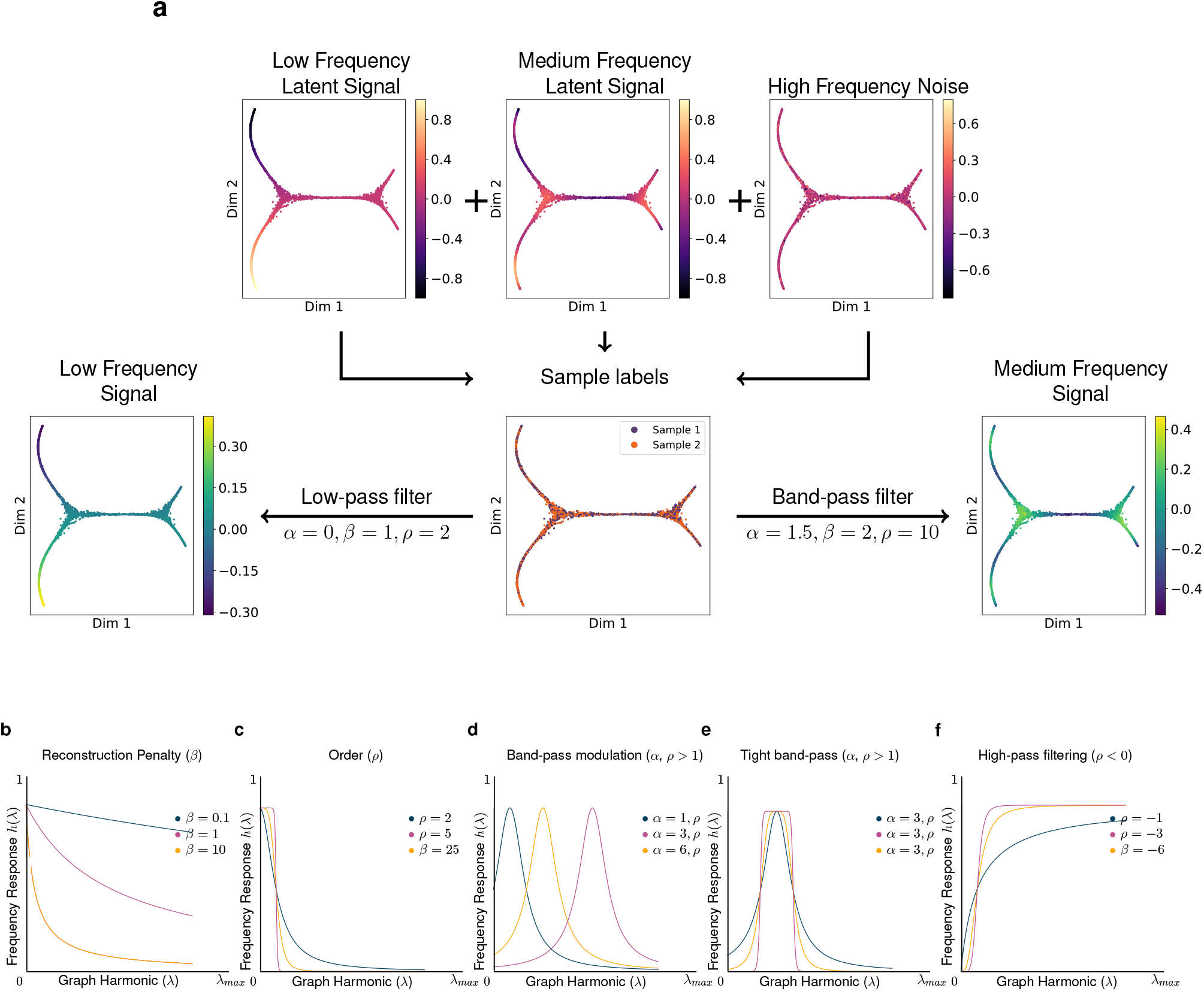
Source Separation and Parameter Analysis with the MELD filter. (**a**) Sample labels (center) are obtained that are a binarized observation of a low frequency latent signal (top left), a medium frequency latent signal (top middle), and high frequency noise (top right). Analysis of the sample labels alone is intractable as they are corrupted by noise and experimental binarization. MELD low-pass filters (bottom left) to separate a longitudinal trajectory and band-pass filters (bottom right) to yield the periodic signature of the medium frequency latent signal. Parameters used for this analysis are supplied beneath the corresponding arrows and the laplacian filter is used for illustrative purposes. (**b**) Reconstruction penalty *β* controls a low-pass filter. For this demonstration, *α* = 0, *ρ* = 1. This filter is equivalent to Laplacian regularization. (**c**) Order *ρ* controls the filter squareness. This parameter is used in the low-pass filter of (**a**). For this demonstration, *β* = 1, *α* = 0. (**d**) Band-pass modulation via *α*. When p is even valued, *a* modulates the central frequency of a band-pass filter. This parameter is used in (**a**) to separate a medium-frequency source from a low-frequency source. (**e**) *α* and *ρ* combine to make square band-pass filters. For (**d**) and (**e**), *β* = 1. (**f**) Negative values of p yield a high-pass filter. For (**b-f**), Laplacian harmonics for a general normalized Laplacian are plotted on the x-axis. The frequency response of the filter given by the colored parameters is on the y-axis.

**Figure S14:**
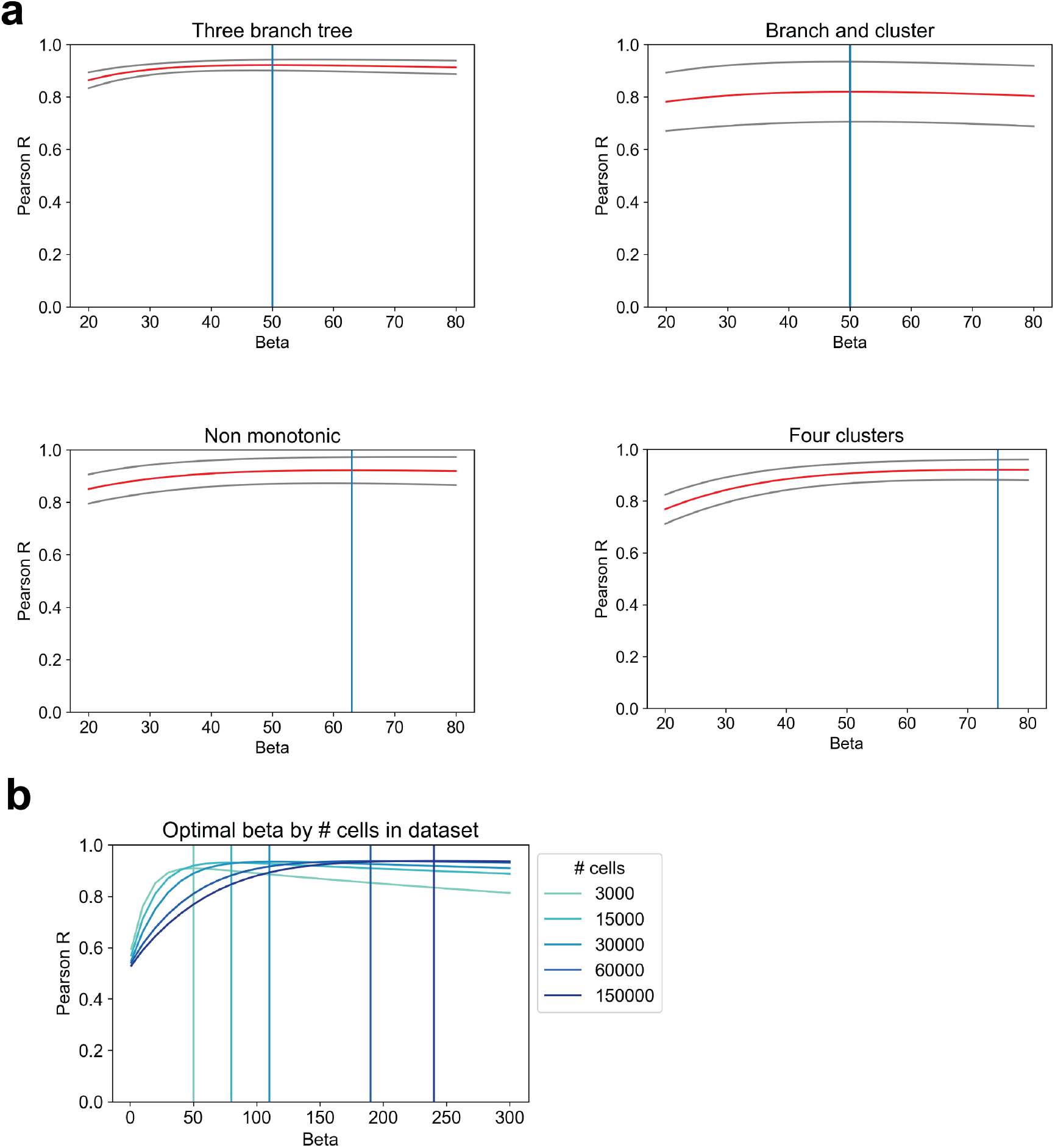
Selecting parameters for MELD. (**a**) Results of a parameter search over the *β* parameter using the four datasets described in Section 4.7. The red line shows the average performance over 10 different datasets of each geometry with one standard deviation marked by the grey lines. We observe reasonably consistent performance of the sample-associated relative likelihood algorithm across all datasets using a *β* value between 50-75. We chose a value of 60 as the default in the MELD package and used this setting for all experiments. (**b**) We observe that the optimal *β* parameter for a dataset varies with the number of cells in the dataset. We suggest increasing the default beta parameter for datasets larger than 30,000 cells.

## Footnotes

1 Freely available at `colab.research.google.com`, most instances provide a 4-core 2GHz CPU and 20GB of RAM.

2 Abbreviations: MLP: Lateral plate, TPM: Tailbud – Presomitic mesoderm, HG: Hatching gland, MBI: Blood island, EPP: Epidermal – pfn1, MEN: Endothelial, PRD: Periderm, EPA: Epidermal anterior, EPO: Otic placode, LLP: Lateral line, EPF: Epidermal – foxi3a, GL: Germline, NRB: Rohon beard, NFP: Floorplate, MHF: Heart field, MPA: Pharyngeal arch, NCC: Neural crest – crestin, END: Endoderm, TSC: Tailbud – spinal cord, NC: Neural crest, NTE: Telencephalon, MPD: Pronephric duct, NHB: Hindbrain, NMB: Midbrain, NTC: Notocord, NDI: Diencephalon, DN: Neurons, OP: Optic

3 Note that in this discussion we abuse notation by treating A as an ordered set of Laplacian eigenvalues and as the diagonal matrix with entries from the elements of this set. Similarly, is both the set of column eigenvectors 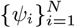 as well as the *N × N* matrix [*ω*_1_ *ω*_2_ … *ω*_N_] with eigenvector as a column.

## Notes

https://github.com/krishnaswamylab/MELD

### Competing Interest Statement

The authors have declared no competing interest.

### Summary of Updates

The paper terminology has been updated significantly. The manuscript has been substantially shortened.

